# A post-transcriptional program of chemoresistance by AU-rich elements and TTP

**DOI:** 10.1101/418715

**Authors:** Sooncheol Lee, Douglas Micalizzi, Samuel S Truesdell, Syed IA Bukhari, Myriam Boukhali, Jennifer Lombardi-Story, Yasutaka Kato, Min-Kyung Choo, Ipsita Dey-Guha, Benjamin T. Nicholson, David T. Myers, Dongjun Lee, Maria A Mazzola, Radhika Raheja, Adam Langenbucher, Nicholas J. Haradhvala, Michael Lawrence, Roopali Gandhi, Christopher Tiedje, Manuel Diaz-Munoz, David A Sweetser, David Sykes, Wilhelm Haas, Daniel A. Haber, Shyamala Maheswaran, Shobha Vasudevan

## Abstract

**Background:** Quiescence (G0) is a transient, cell cycle-arrested state. By entering G0, cancer cells survive unfavorable conditions such as chemotherapy and cause relapse. While G0 cells have been studied at the transcriptome level, how post-transcriptional regulation contributes to their chemoresistance remains unknown.

**Results:** We induced chemoresistant and quiescent (G0) leukemic cells by serum-starvation or chemotherapy treatment. To study post-transcriptional regulation in G0 leukemic cells, we systematically analyzed their transcriptome, translatome, and proteome. We find that our resistant G0 cells recapitulate gene expression profiles of *in vivo* chemoresistant leukemic and G0 models. In G0 cells, canonical translation initiation is inhibited; yet we find that inflammatory genes are highly translated, indicating alternative post-transcriptional regulation. Importantly, AU-rich elements (AREs) are significantly enriched in the up-regulated G0 translatome and transcriptome. Mechanistically, we find the stress-responsive p38 MAPK-MK2 signaling pathway stabilizes ARE mRNAs by phosphorylation and inactivation of mRNA decay factor, tristetraprolin (TTP) in G0. This permits expression of ARE-bearing TNFα and DUSP1 that promote chemoresistance. Conversely, inhibition of TTP phophorylation by p38 MAPK inhibitors and non-phosphorylatable TTP mutant decreases ARE mRNAs and sensitizes leukemic cells to chemotherapy. Furthermore, co-inhibiting p38 MAPK and TNFα—prior to or along with chemotherapy—substantially reduced chemoresistance in primary leukemic cells *ex vivo* and *in vivo*.

**Conclusions:** These studies uncover post-transcriptional regulation underlying chemoresistance in leukemia. Our data reveal the p38 MAPK-MK2-TTP axis as a key regulator of expression of ARE bearing mRNAs that promote chemoresistance. By disrupting this pathway, we developed an effective combination therapy against chemosurvival.

## Background

Quiescent (G0) cells are an assortment of reversibly arrested cells, including dormant stem cells, which are found as a clinically relevant subpopulation in cancers (1–4). Such cells are anti-proliferative, anti-differentiation, and anti-apoptotic, and show distinct properties including resistance to harsh conditions (1;2;5–10). G0 cells show specific gene expression that may underlie their resistance and other properties (1;2;8–10). Analyses from multiple groups revealed some genes up-regulated at the transcriptional level (1;8;11). Altered polyadenylation site selection on mRNAs produces longer 3□-untranslated regions (3□UTRs) in G0 compared to proliferating cells— which increases 3□UTR elements that can mediate post-transcriptional gene expression regulation (12). Our previous data demonstrated that translation mechanisms are distinct in G0 leukemic cells, with decreased canonical translation mechanisms and increase in mRNA translation by alternative mechanisms that involve non-canonical translation initiation factors (13) and 3’UTR mediated specific mRNA translation (14). These data suggest that alternate post-transcriptional mechanisms in G0 cancer cells may regulate a distinct translatome to mediate their resistance. Translated genes, post-transcriptional mechanisms involved, and outcomes on cancer persistence remain to be investigated.

We analyzed the translatome and proteome of chemotherapy-surviving G0 cancer cells, focusing on acute monocytic leukemia (AML), to provide comprehensive information that complement and expand previous transcriptome analyses (1;2;8;11;15;16), revealing critical genes that are post-transcriptionally regulated for chemo-survival. G0 can be induced by growth factor-deprivation or serum-starvation and other conditions that isolate dormant cancer stem cells in distinct cell types (1;6;7). Our data demonstrate that serum-starvation induced G0 AML cells are chemoresistant—similar to surviving AML cells, isolated after chemotherapy. Chemoresistant cells isolated via serum-starvation, or as surviving cells post-chemotherapy, show inhibition of canonical translation mechanisms, indicating that non-canonical mechanisms express specific mRNAs when these cells are chemoresistant. Consistently, the translatomes and proteomes of serum-starved G0 and chemo-surviving cells show greater similarity than the transcriptome alone. Our data reveal that DNA damage and stress signaling cause post-transcriptional alterations to produce a specialized gene expression program of pro-inflammatory, immune effectors that elicit chemosurvival.

## Results

### Serum-starvation or AraC treatment induces a quiescent and chemoresistant state of leukemic cells

To study clinical resistance in cancer, THP1 human AML cells were used as they show significant resistance to AraC (17) (cytosine arabinoside, Fig. S1A), a standard anti-leukemic chemotherapeutic that targets DNA replication and thus proliferating cells (referred to as S+). Our data and others find that serum-starvation of THP1 (13) and other cell lines (1;8;11;18) induces a transient G0 state with known G0 and cell cycle arrest markers expressed (Fig. 1C-D, S1B-C). Such serum-starvation induced G0 cells (referred to as SS) can be returned to the cell cycle upon serum addition (Fig. 1D), verifying that they are quiescent and transiently arrested, unlike senescence or differentiation that are not easily reversed. We find that serum-starvation induced G0 SS cells show resistance to AraC chemotherapy. Serum-grown S+ cells show a dose-dependent decrease in cell viability with AraC as expected, while SS cells persist, indicating their chemoresistance (Fig. 1E). Chemoresistant cancer cells include cancer stem cells and are a subpopulation that can be isolated from cancers after treatment with chemotherapy (2;6–10) that targets and eliminates S+ cells. We find that AraC-surviving THP1 (referred to as AraCS) cells are transiently arrested, like SS cells (Fig. 1C-D, S1B); both AraCS and SS cells survive chemotherapy (Fig. 1E). AraCS cells recover from their transient arrest upon AraC removal, proliferate (Fig. 1D), affirming the reversible G0 arrest state of chemoresistant cells, similar to SS cells (1;2;6–10).

**Figure 1.**
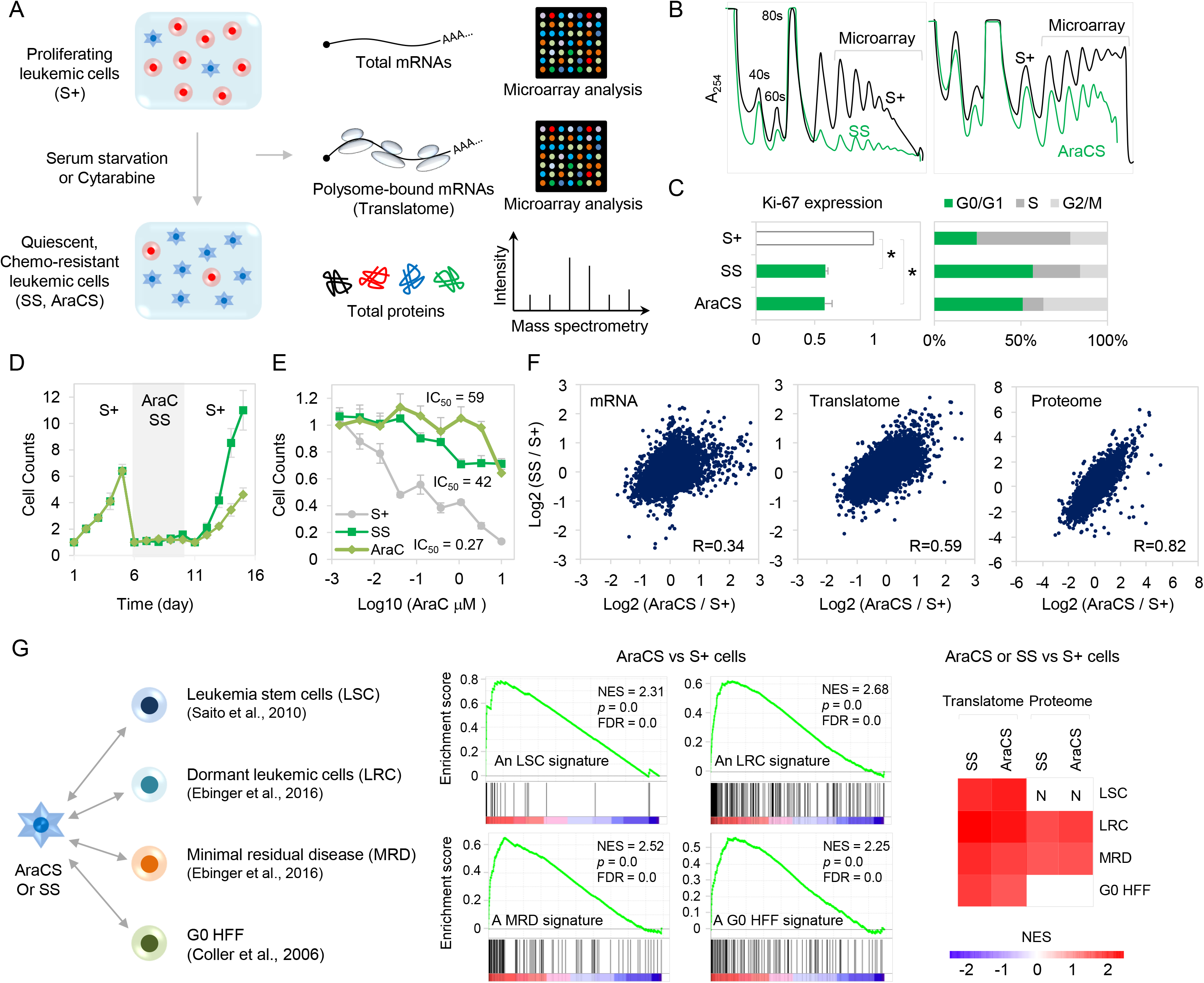
G0 leukemic cells induced by AraC or serum-starvation are chemoresistant and recapitulate gene expression programs of *in vivo* chemoresistant and G0 models. **A**. Transcriptome, translatome and proteome analyses in proliferating and G0 leukemic cells. G0 cells (AraCS, SS cells) were induced by treatment of proliferating cells (S+) with AraC or serum starvation. Total RNAs, polysome-associated mRNAs and protein were analyzed by comparative microarray and quantitative proteomics. **B**. Polysome profiles of S+, SS and AraCS are shown. Polysome-associated mRNAs were isolated and analyzed by microarray. **C**. Ki67 translatome level and flow cytometric quantification of G0/G1, S and G2/M phases, using BrdU and PI staining. **D**. Cell counting with trypan blue staining. Proliferating THP1 cells were serum-starved or treated with AraC for days specified. Then, serum was added to SS cells while AraCS cells were resuspended in fresh media. **E**. S+, SS and AraCS cells were treated with various concentration of AraC for 3 days. Viable THP1 leukemic cells were measured by cell counting using trypan blue staining and IC_50_ values of AraC are shown. **F**. Comparison of transcriptomic, translatomic and proteomic changes in response to SS and 5 μM AraC treatments. **G**. Comparison of AraCS and SS with leukemic stem cells (LSC) (16) in AML, dormant leukemic cells (LRC) (15), minimal residual disease (MRD) (15) in ALL, and G0 fibroblasts (1). GSEA analysis was performed to determine whether previously published transcriptome signatures of LSC, LRC, MRD and G0 HFF are up-regulated in AraCS and SS cells, compared to S+ cells. ‘N’ marks the limited resolution of the proteome in the GSEA. \**P*□≤□0.05. Data are represented as average ± SEM. See also Fig. S1 & Table S1.

### G0 cells induced by SS or AraC have similar translatomes and proteome features that recapitulate gene expression profiles of *in vivo* chemoresistant leukemic and G0 models

To study post-transcriptionally regulated genes in G0, we profiled S+, SS cells and AraCS cells at the proteome, translatome and transcriptome levels using multiplexed quantitative proteomics (14), microarray analysis of heavy polysome-associated mRNAs (13;14;19), and total RNAs respectively (Fig. 1A-B, S1D-F). Notably, we find that AraCS and SS cells show more similar gene expression profiles at the proteome and translatome levels, compared to transcriptome levels (Fig. 1F). These data suggest that although these chemoresistant G0 cells are isolated *via* two different methods, they exhibit a common set of translatome and proteome, which could underlie their common characteristic of chemoresistance. These data indicate the relevance of examining both the translatome and transcriptome. Time-course translatome analysis revealed that SS G0 cells that were serum-starved for short periods (4 hours and 1 day), are distinct from SS G0 cells that were serum-starved for long periods (2 days and 4 days) (Fig. S1E-F). This is consistent with G0 as a continuum of assorted, arrested states (1), with temporal differences in underlying gene expression in early G0 compared to more homogeneity at late G0. SS and AraCS cells provide sufficient material to perform concurrent translatome, proteome and transcriptome profiling, compared to limited cells from in *vivo* resistance models where only transcriptomes were profiled. To test whether our G0 leukemic cells are relevant models to study chemoresistance and G0, gene expression profiles of AraCS and SS cells were compared to published transcriptome profiles of leukemia stem cells (LSC) from AML (16), dormant leukemic cells (LRC), and minimal residual disease (MRD) from chemotherapy surviving patient samples with acute lymphocytic leukemia (ALL) (15), as well as SS G0 fibroblasts (G0 HFF) (1). Importantly, we find that these published transcriptome signatures for *in vivo* chemoresistance and G0 models were significantly up-regulated in our SS and AraCS cells (referred to as resistant G0 leukemic cells), compared to S+ cells (Fig. 1G, S1G). These data indicate that our resistant G0 leukemic cells are relevant models to study post-transcriptional regulation in chemoresistance as they have similar gene expression profiles to known transcriptional profiles from *in vivo* chemoresistance models.

### Inhibition of canonical translation initiation in resistant G0 leukemic cells

We find overall protein synthesis is reduced at least 2-fold in AraCS, compared to S+ cells (Fig. 2B, S1D). Mechanistically, both rate-limiting steps in canonical translation initiation: recruitment of initiator tRNA, and mRNA cap recognition to recruit mRNAs to ribosomes are inhibited in G0 leukemic cells (Fig. 2A). Recruitment of initiator tRNA by eIF2 can be blocked by eIF2α phosphorylation as a stress response (13;20–25). We find that two eIF2 kinases, PKR and PERK, are activated and increase eIF2α phosphorylation (Fig. 2C) in G0 leukemic cells to inhibit canonical translation initiation. Consistent with our previous study (14), we observed dephosphorylation of 4E-BP (Fig. 2C) that inhibits cap-dependent translation initiation (26;27). Low mTOR activity is known to reduce translation of terminal oligopyrimidine tract (TOP) mRNAs such as ribosomal protein mRNAs (26;28;29), which is decreased in SS and AraCS cells (Fig. 2D). Decreased canonical translation can enable post-transcriptional regulation of specific genes, as observed previously (13;14) and lead to survival of G0 leukemic cells.

**Figure 2.**
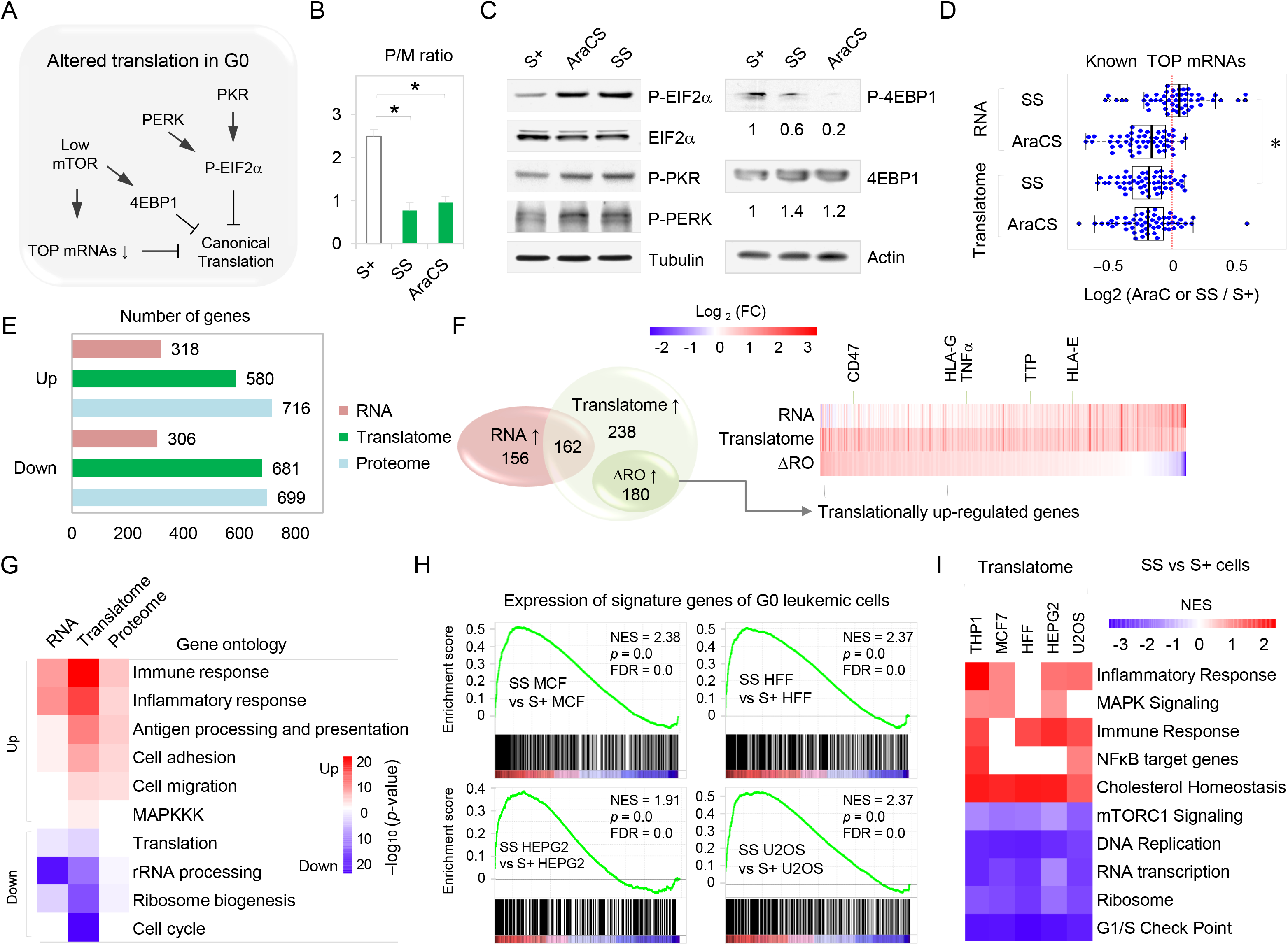
Inflammatory response mRNAs are selectively translated in G0 leukemic cells, where canonical translation is inhibited. **A**. Repression of canonical translation. **B**. Polysome to monosome ratios in S+, SS and AraCS. **C**. Western analysis of translation initiation factor, eIF2α and its regulators eIF4EBP, PERK and PKR. **D**. Boxplot of the transcriptome and translatome changes in known TOP mRNAs in response to SS or AraC treatment. **E**. Number of differentially expressed genes. **F**. Venn diagram shows 162 genes up-regulated at both the transcriptome and translatome levels and 180 genes translationally up-regulated, where ribosome occupancy, RO increases by at least 1.5-fold. Heatmap shows gene expression changes in RNA, translatome levels and RO. **G**. Gene ontology (GO) analyses of differentially expressed genes shown in Fig. 2E. Statistical significance of enriched GO categories is shown as a heatmap. **H**. Expression of signature genes of G0 leukemic cells in published transcriptomes of in vivo resistant leukemic and G0 models in other G0 cells. **I**. Translatome analysis of G0 cells from five different cell types. Heatmap of normalized enrichment score (NES) is shown. \**P*□≤□0.05. Data are represented as average ± SEM. See also Fig. S2 & Table S1.

### Global translatome analysis shows that inflammatory response genes are selectively translated in resistant G0 cancer cells

We measured the number of genes upregulated at the transcriptome, translatome and proteome levels in resistant G0 leukemic cells, compared to S+ cells. A significantly greater number of genes were upregulated in the translatome (580 genes, Table S1) and proteome (716 genes), compared to the transcriptome (318 genes) as shown in Fig. 2E. Importantly, 57% of upregulated genes were upregulated only in the translatome level (Fig. 2F) but not in the transcriptome, indicating post-transcriptional regulation. To investigate the biological function of these differentially expressed genes, gene ontology (GO) analysis was performed. Gene categories up-regulated in G0 translatomes include inflammatory response, immune modulators (receptors, antigen presentation and processing genes), cell adhesion, cell migration, lipid biosynthesis and cholesterol pathway genes (Fig. 2G, S2E). Down-regulated genes include RNA processing and ribosome genes (Fig. 2G). To identify translationally up-regulated genes, we measured the change in ribosome occupancy (RO) which is the ratio of polysome-associated mRNA levels to total mRNA levels of each gene (Fig. 2F, Venn diagram, heat map). We find 180 genes are translationally up-regulated above RNA level changes. These genes include antigen processing and presentation genes (30) (HLA-G, HLA-E) and immune receptors (CD47, Fig. 2F-G, S2I) (31–33) that regulate anti-tumor immune response and are associated with leukemic stem cells and resistance (34;35).

We asked if this specific gene expression profile in resistant G0 leukemic cells is conserved in G0 cells of other tumors and normal cells. Therefore, global translatome profiling was conducted in G0 cells from four different cells lines: breast cancer (MCF7), liver cancer (HEP-G2), and osteosarcoma (U2OS) as well as non-cancerous fibroblasts (HFF) (Fig. S2). Their translatome profiles were compared with resistant G0 leukemic cells, using GSEA and DAVID tools (Fig. 2H-I, S2E-F). We find that 580 signature genes of resistant G0 leukemic cells (Table S1) were highly upregulated at the translatome level in G0 cells of these other cell types (Fig. 2H). As expected for these arrested cells, genes related to cell cycle, ribosome biogenesis, and DNA replication were commonly down-regulated (Fig. 2I, S2E). Importantly, inflammatory response genes were commonly up-regulated in cancer G0 cells but not normal G0 fibroblasts and do not significantly overlap with senescence-associated secretory pathway (SASP) (Fig. 2I, S2G) (36;37), indicating a specific role in chemoresistant cancer cells.

### Stabilization of ARE-bearing mRNAs is mediated by phosphorylation of TTP in resistant G0 leukemic cells

To identify *cis*-acting elements that mediate post-transcriptional regulation, the untranslated regions (UTRs) of differentially expressed genes were examined. We find that a GC-rich motif was enriched on 5’UTRs of translationally up-regulated genes and an AU-rich motif, on 5’UTRs of down-regulated genes, indicating that mRNAs with structured 5’UTRs are highly translated in G0 cells (Fig. S3A-B). Importantly, 3’UTR AU-rich elements (AREs) are significantly enriched in the up-regulated translatome as well as transcriptome (Fig 3A). Furthermore, 30% of the translatome signature of G0 leukemic cells bear AREs (Table S2), including proinflammatory cytokines such as TNFα and chemokines (Fig. 3B-C). AREs are important post-transcriptional regulatory elements that mediate rapid degradation and repression of mRNAs (38). To understand how ARE mRNAs are highly expressed in G0 cells, we assessed the expression level of RNA-binding proteins. As expected, ARE-binding proteins known to cause mRNA decay or translation repression (39;40) are significantly reduced in G0 cells (Fig. S3F). Additionally, the exosome and proteasome complexes that are implicated in ARE mRNA decay (41) (42) are reduced (Fig. S3C-E). However, a key ARE mRNA decay and translation repression factor, Tristetraprolin (TTP) was surprisingly increased in AraCS from multiple AML cell lines (Fig. 3D-E). However, we find that TTP is phosphorylated in SS and AraCS cells (Fig. 3E, right). TTP phosphorylation is established to increase its levels (43), and block its ability to destabilize ARE mRNAs, thus enabling ARE mRNA translation upon LPS treatment (44;45). To test whether phosphorylation of TTP was required for the increased expression of ARE mRNAs in G0 leukemic cells, we generated non-phosphorylatable mutant TTP with key phosphorylation sites (Ser 52, 178) replaced by alanine (TTP-AA). TTP-AA has been shown to dominantly maintain ARE mRNA decay activity and reduce pro-inflammatory cytokines like TNFα in immune cells (43–45). Expression of myc-tagged TTP-AA significantly reduced TNFα mRNA in both THP1 and K562 AraCS cells (Fig. 3F), overturning the decay inactivity of endogenous phospho-TTP. To determine the effect of TTP phosphorylation on the stability of ARE mRNAs, we measured the half-life of TNFα mRNA. Expression of TTP-AA mutant more significantly reduced the half-life of TNFα mRNA than TTP wild-type expressed in AraC-treated TTP-deficient macrophages ^48^ (Fig 3G). Furthermore, immunoprecipitation demonstrated that TTP-AA was associated with TNFα mRNA in AraCS cells (Fig 3H). These data indicate that inactivation of ARE mRNA decay by TTP phosphorylation (43;45;46) is a key regulator of expression of a pro-inflammatory gene, TNFα, in chemoresistant G0 cells. These results are consistent with our findings of increased levels and translation of ARE bearing mRNAs due to decreased ARE mRNA decay activity in G0 cells (Fig. 3A-C, S3C-F).

**Figure 3.**
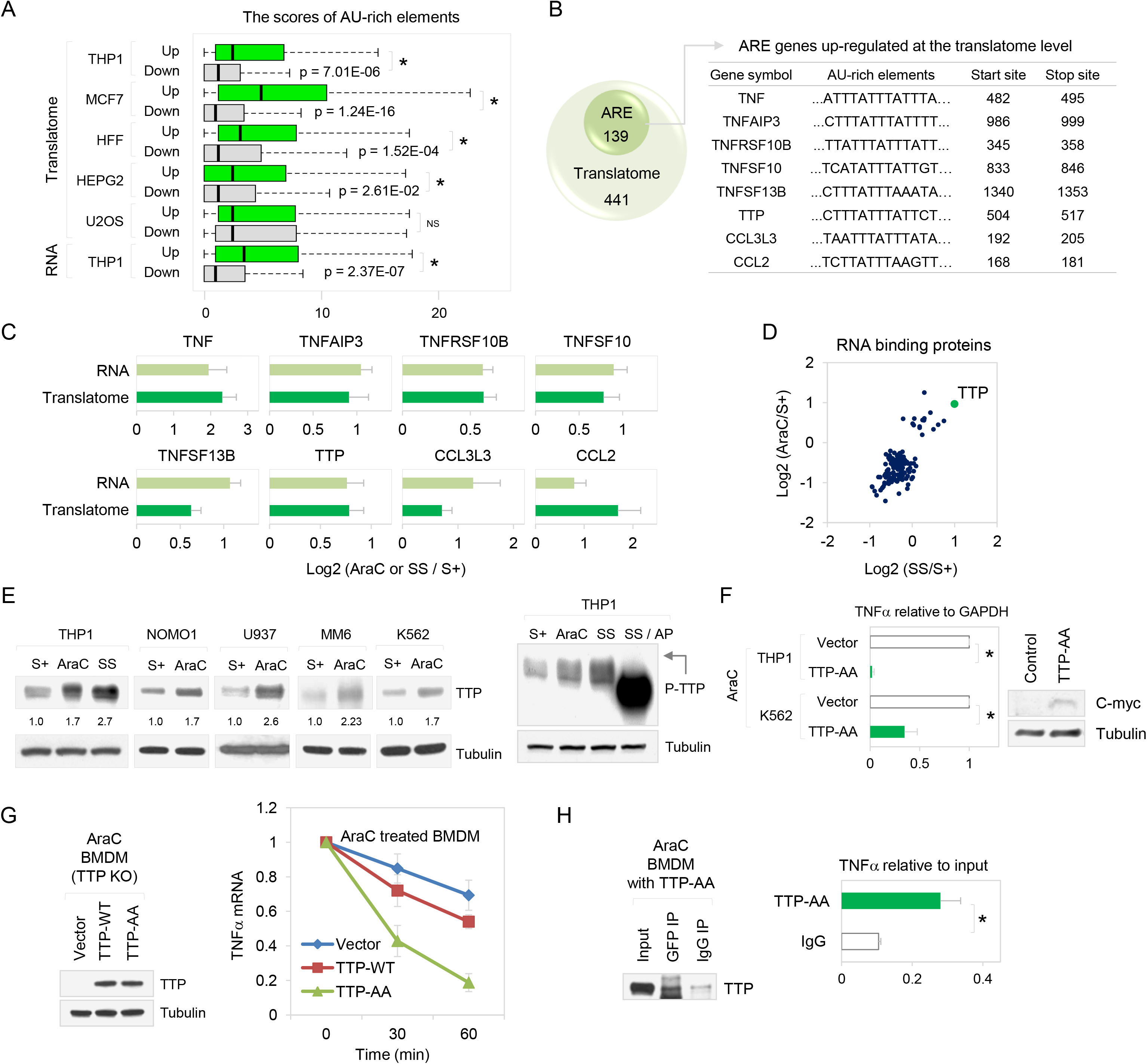
Phosphorylation of TTP stabilizes ARE-bearing TNFα in G0 leukemic cells. **A**. Boxplot of ARE scores (SI methods) in the 3’UTRs of genes which are up- or down-regulated at the translatome or RNA levels in G0 compared to S+ cells. **B**. Venn diagram shows genes that are up-regulated at the translatome level and contain AREs (left). List of such genes (right, Table S2). **C**. Expression of ARE genes at the RNA and translatome levels. **D**. Scatter plot showing the expression of RNA binding protein genes from RBPDB database (SI methods). TTP is indicated with a green dot. **E**. Western analysis of TTP in lysates from multiple leukemic cell lines in the absence or presence of alkaline phosphatase (AP). Phospho-TTP is indicated with an arrow. **F**. Bar graph shows TNFα mRNA expression normalized to GAPDH mRNA upon over-expression of vector or c-myc tagged non-phosphorylatable mutant TTP (TTP-AA) in AraC-treated THP1 or K562 cells. Western analysis of TTP-AA with c-myc antibody (right). **G**. Half-life of TNFα mRNA. TTP-deficient BMDM cells were transduced with doxycycline inducible plasmids that express GFP vector, TTP wild-type or TTP-AA mutant. Cells were induced with 1 μg/ml doxycycline prior to 1 μM AraC treatment. Western analysis of induction of TTP protein. TNFα mRNA level was measured at indicated time points by qPCR after transcriptional arrest with 5 μg/ml actinomycin D treatment. **H**. Association of TTP-AA with TNFα mRNA in AraCS cells. TTP-AA was immunoprecipitated with GFP antibody from AraC-treated BMDM cells expressing GFP-tagged TTP-AA. (Western blot), followed by qPCR analysis of TNFα mRNA (graph). \**P*□≤□0.05. Data are represented as average ± SEM. See also Fig. S3 & Table S2.

### The p38 MAPK-MK2 pathway phosphorylates TTP to promote expression of ARE-bearing mRNAs in resistant G0 leukemic cells

To investigate how TTP is phosphorylated in resistant G0 leukemic cells, we examined key signaling molecules involved in DNA-damage response (DDR) (Fig. 4A) that is induced by chemotherapies like AraC (47–50). As expected, AraC treatment induced rapid phosphorylation and activation of ATM (Fig. 4B). Importantly, we find that these conditions lead to phosphorylation and activation of p38 MAPK and its downstream effector, MAPKAPK2 (MK2) (51;52) (Fig. 4B). MK2 has been shown to phosphorylate TTP in macrophages treated with lipopolysaccharide (LPS) (43;45;46). To examine whether the p38 MAPK-MK2 pathway phosphorylates TTP in resistant G0 leukemic cells, two different inhibitors of p38 MAPK were tested. Treatment with p38 MAPKα/β inhibitor, LY2228820 (LY) (52;53), or a pan-p38 MAPK inhibitor that targets all isoforms, BIRB796 (BIRB) (54), blocked phosphorylation of MK2 and prevented MK2-mediated TTP phosphorylation and reduces TNFα in AraCS cells (Fig. 4C). These results suggest that p38 MAPK-MK2 phosphorylates TTP, resulting in enhanced expression of ARE mRNAs such as TNFα upon AraC treatment (Fig. 4A). To test if the p38 MAPK-MK2-TTP pathway regulates TNFα expression via its ARE, a firefly luciferase reporter bearing the 3’ UTR ARE of TNFα, and as control, Renilla luciferase, were co-transfected. Luciferase activity of the ARE reporter increased by 2-fold in AraCS cells compared to S+ cells but not when p38 MAPK was inhibited (Fig. 4D). These data suggest that the p38 MAPK-MK2-TTP axis up-regulates expression of specific genes via AREs in G0 leukemic cells.

**Figure 4.**
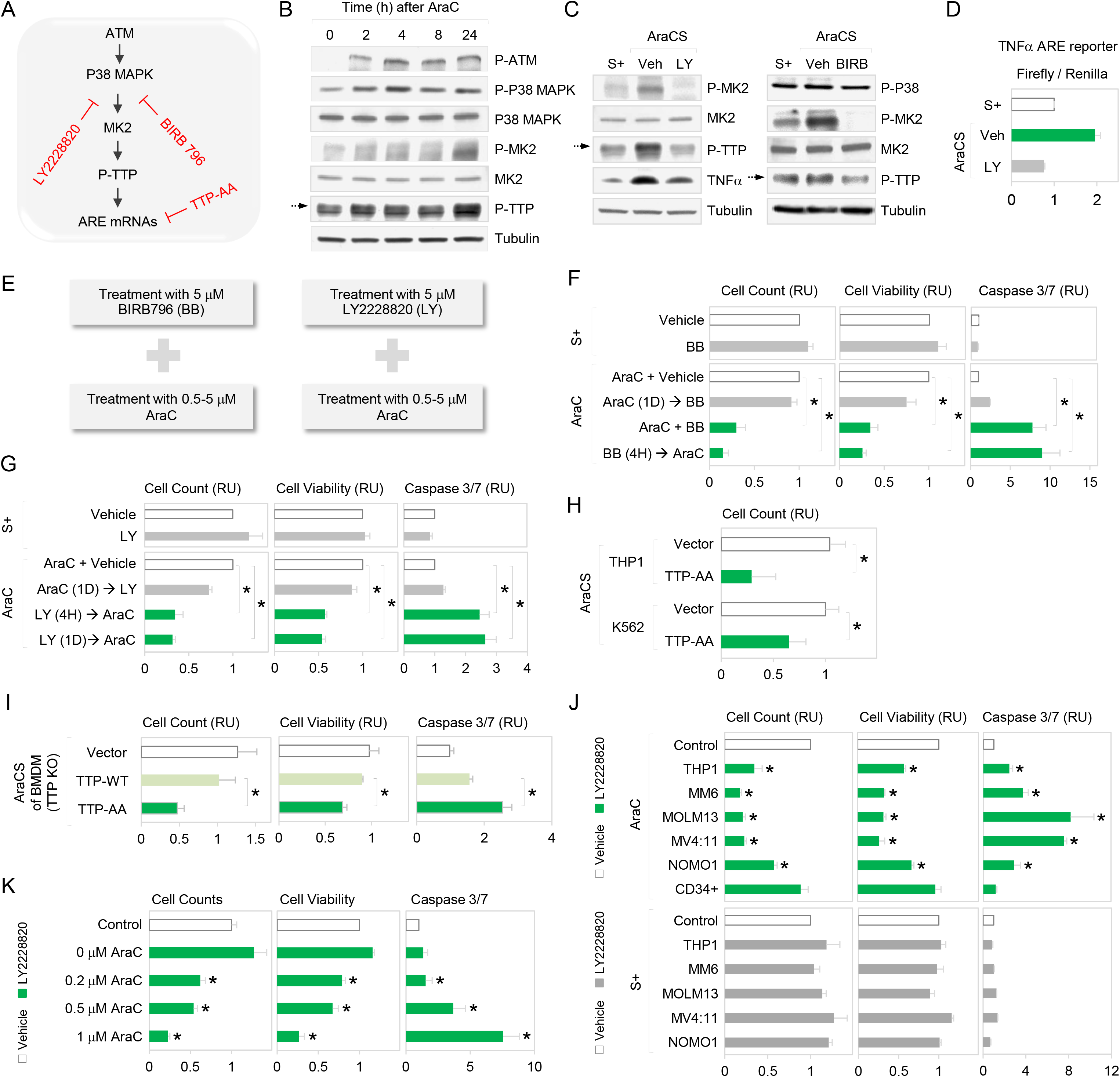
Phosphorylation of TTP by p38 MAPK-MK2 promotes chemoresistance. **A**. The p38 MAPK (p38)-MK2 pathway enables stabilization and translation of ARE-bearing mRNAs via TTP phosphorylation in chemoresistant G0 cells. LY2228820 (LY) and BIRB396 (BB) are p38 inhibitors. **B**. Western analysis of in lysates from THP1 cells at indicated time points after AraC treatment. **C**. Western analysis in S+ and AraCS cells treated with vehicle, 5 μM LY or 5 μM BB. **D**. Firefly luciferase activity of a reporter bearing TNFα ARE in its 3’UTR normalized to activity of co-transfected Renilla luciferase in S+ and AraCS cells treated with vehicle, 5 μM LY. **E**. Sequential treatment with p38 inhibitors and AraC in leukemic cells. **F-G**. Effect of p38 inhibitions on survival of AraC-resistant cells. THP1 cells were treated with 5 μM BB, 5 μM LY, and vehicle in the absence (S+, top panels) or presence (AraC, bottom panels) of 5 μM AraC treatment for three days. Bar graphs show relative cell viability and death assessed by cell counting, MTS and caspase 3/7 assays. In the presence of AraC, THP1 cells were treated with p38 inhibitors prior to AraC treatment (BB → AraC, LY → AraC), at the same time with AraC (AraC + BB) and 1 day after AraC (AraC → BB, AraC → LY). 4H and 1D indicate 4 hours and 1 days, respectively. **H-I**. Effect of TTP-AA mutant on survival of AraC resistant cells. TTP-AA mutant expression prior to 5 μM AraC treatment, decreased TNFα in THP1 or K562 cells (Fig. 3F). Cell viability was assessed by cell count (H). TTP-AA, TTP wild-type and vector were expressed in TTP-deficient BMDM cells prior to 1 μM AraC treatment. Bar graphs show relative cell viability and death (I). **J**. Effect of p38 inhibition on resistant cells from five AML cell lines (M5 FAB subtype). Cells were treated with 5 μM LY or vehicle 4 hours prior to AraC treatment (top panel, AraC) or in the absence of AraC (bottom panel, S+). Human CD34+ cells from healthy donors were tested as a control. **K**. Effect of p38 inhibition on survival of chemoresistant cells induced with various concentrations of AraC. MV4:11 leukemic cells were treated with 5 μM LY or vehicle prior to 0 μM, 0.2 μM, 0.5 μM or 1 μM AraC for 3 days. \**P*□≤□0.05. Data are represented as average ± SEM. See also Fig. S4.

### Phosphorylation of TTP induced by p38 MAPK-MK2 promotes chemoresistance

We noted that the p38 MAPK-MK2 pathway was rapidly activated to phosphorylate TTP within one day of SS or AraC treatment (Fig. 4B, S4A-B). To test the effect of inhibition of TTP phosphorylation on chemoresistance, p38 MAPK was inhibited *before* (or along with) as well as *after* treatment with AraC— and then chemosurvival was measured using multiple assays, including cell death and two cell viability assays (Fig. 4E-G). Inhibition of p38 MAPK with BIRB or LY, one day *after* AraC treatment, when TTP was already phosphorylated, did not show any significant reduction in survival of AraC-resistant cells (Fig. 4F-G). Conversely, inhibition of p38 MAPK at earlier time points prior to AraC treatment, when TTP was not phosphorylated, increased apoptosis and reduced survival of AraC-resistant cells (Fig. 4F-G). As a control, p38 MAPK inhibition alone does not affect viability of S+ cells that are not treated with AraC (Fig. 4F-G). These results suggest that p38 MAPK is rapidly activated upon AraC treatment to turn on downstream survival pathways such as phosphorylation of TTP. Thus, to inhibit phosphorylation of TTP and hence overcome AraC resistance effectively, p38 MAPK needs to be targeted at early time points.

To confirm that phosphorylation of TTP induces chemoresistance, we over-expressed TTP mutant (TTP-AA) that cannot be phosphorylated by p38 MAPK-MK2, followed by AraC treatment. Importantly, we find that TTP-AA mutant expression reduces survival of AraC-resistant cells in THP1 and K562 leukemic cell lines (Fig. 4H). Furthermore, TTP-AA mutant, expressed in TTP-knockout macrophages, induced apoptosis of AraC-surviving cells more significantly, compared to TTP wild-type (Fig. 4I). Consistently, in multiple AML cell lines, early inhibition of p38 MAPK showed dramatically reduced chemosurvival but not in non-cancerous CD34+ cells (Fig. 4J). When treated with p38 MAPK inhibitor alone, viability of S+ cells in multiple AML cell lines remained unchanged, indicating the synergism of AraC and p38 MAPK inhibitors (Fig. 4J). Interestingly, p38 MAPK inhibition eliminated resistant cells more significantly at increasing concentrations of AraC (Fig. 4K). This indicates that treatment with high concentrations of AraC would increase the number of cells induced into the resistant G0 state with strong phosphorylation of p38 MAPK-MK2-TTP. Conversely, even low concentrations of BIRB were sufficient to reduce chemoresistance (Fig. S4C). Unlike in solid tumors, where activation of p38 MAPK-MK2 induces resistance by arresting the cell cycle (38;51;52), p38 MAPK inhibition did not affect the cell cycle in AML cells (Fig. S4D). These data uncover rapid activation of a p38 MAPK-MK2 pathway that enables chemosurvival of G0 leukemic cells via inhibition of TTP activity.

### TNFα, induced by phosphorylation of TTP, promotes chemoresistance

We demonstrated that TTP regulates the stability of ARE mRNAs such as TNFα in AraCS cells (Fig. 3G). Furthermore, inactivation of TTP allowed elevated TNFα translatome and protein levels in resistant G0 leukemic cells (Fig 5B-C). To assess the effect of TNFα on chemoresistance, we altered TNFα levels genetically and phamacologically in G0 cells (Fig. 5A). Induction of TNFα depletion prior to AraC effectively reduced AraC resistance, compared to depleting TNFα after AraC treatment, while no effect was observed with TNFα depletion alone without AraC (Fig. 5D). In contrast, addition of recombinant TNFα enhanced survival of AraCS cells (Fig. 5D). TNFα-mediated chemoresistance is not due to arrested cell cycle as TNFα treatment without subsequent AraC does not alter the cell cycle (Fig. S5C). These data suggest that phosphorylation of TTP and subsequent expression of TNFα, which are induced by p38 MAPK-MK2, are responsible for survival of G0 leukemic cells.

**Figure 5.**
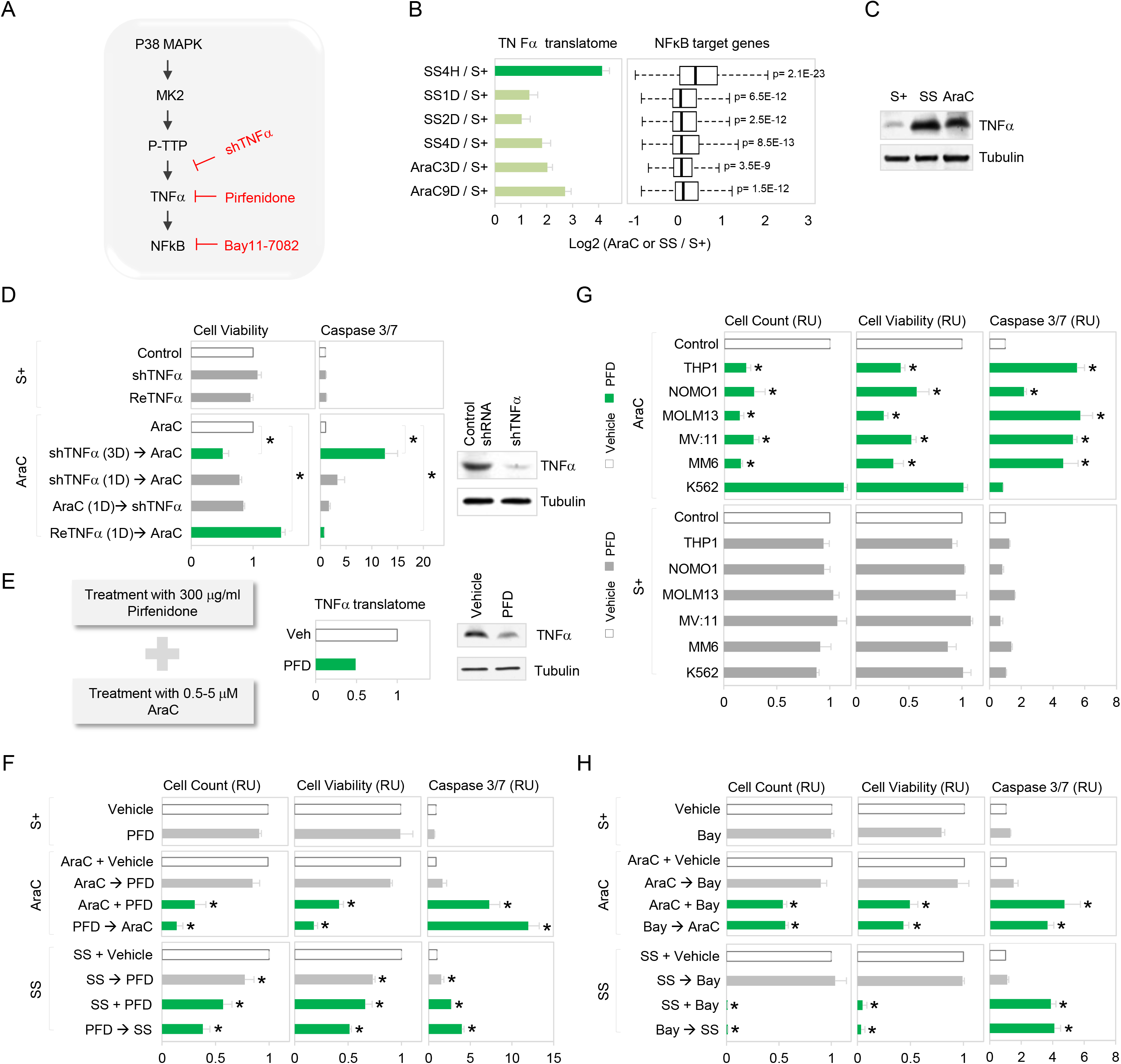
TNFα induced by phosphorylation of TTP promotes chemoresistance. **A**. Phosphorylation of TTP by the p38-MK2 pathway stabilizes ARE-bearing TNFα mRNA, resulting in activation of NF-kB signaling in resistant G0 leukemic cells. TNFα expression is inhibited by TTP-AA mutant, pirfenidone (PFD) or shRNAs, and NF-kB signaling by NF-kB inhibitor, Bay11-7082. **B**. Expression of TNFα and NF-kB target genes at the translatome level at indicated time points after SS or AraC treatment. **C**. TNFα protein level in S+, SS and AraCS cells. **D**. Effect of TNFα on chemoresistance. THP1 cells were transduced with doxycycline inducible shRNA against TNFα or control shRNA. ShRNA against TNFα was induced prior to AraC (shTNFα → AraC) or after AraC (AraC → shTNFα) and recombinant TNFα protein was added 1 day prior to AraC (ReTNFα → AraC). Cell viability and western analysis of TNFα, are shown. **E**. Effect of PFD on TNFα expression at the translatome (middle) and protein levels (right) in AraCS cells. **F**. Effect of pharmacological inhibition of TNFα by PFD on AraC resistance. THP1 cells were treated with 300 μg/ml PFD or vehicle in the absence of AraC (S+, top panels), in the presence of AraC (AraC, middle panels), or on serum starvation (SS, bottom panels). Bar graphs show cell viability and death assessed by cell counting, MTS and caspase 3/7 assays. In middle or bottom panels, THP1 cells were treated with PFD 1 day prior to AraC or SS (PFD → AraC, PFD → SS), at the same time with AraC or SS (AraC + PFD, SS + PFD), and 1 day after AraC or SS (AraC → PFD, SS → PFD). **G**. Effect of TNFα inhibition on AraC resistance from six different leukemic cell lines. Cells were treated with PFD or vehicle 1 day prior to AraC (AraC, top panels) or in the absence of AraC (bottom panels, S+). **H**. Effect of NF-kB inhibition on AraC resistance. THP1 cells were treated with 10 μM Bay11-7082 or vehicle in the absence of AraC (S+, top panels), in the presence of AraC (AraC, middle panels) or under serum starvation (SS, bottom panels). In middle or bottom panels, THP1 cells were treated with Bay11-7082 1 day prior to AraC or SS (Bay → AraC, Bay → SS), at the same time with AraC or SS (AraC + Bay, SS + Bay), and 1 day after AraC or SS (AraC → Bay, SS → Bay). \**P*□≤□0.05. Data are represented as average ± SEM. See also Fig. S5.

TNFα can also be inhibited pharmacologically with the drug pirfenidone (PFD) that can block TNFα translation in RAW264.7 cells and is used to treat idiopathic pulmonary fibrosis (52;55;56). In G0 leukemic cells, PFD reduced TNFα translatome and protein levels but not mRNA levels (Fig. 5E, S5F). PFD treatment at least 18 hours *prior to* or along with AraC or SS significantly reduced viability of G0 leukemic cells but failed to reduce resistance when added *after* AraC treatment (Fig. 5F, S5D). As observed with p38 MAPK-MK2 activation (Fig. 4A-B), TNFα translatome level also is rapidly and dramatically increased upon SS treatment (Fig. 5B). These data indicate that activation of TNFα is an early event in G0 induction, which leads to resistance, and needs to be inhibited early to preclude downstream survival regulators. PFD treatment alone does not affect the viability of untreated S+ cells, indicating that the cytotoxic effect of PFD is specific to G0 leukemic cells (Fig. 5F). PFD treatment reduced chemotherapy survival in multiple AML cell lines (Fig. 5G). Similar results were observed in MCF7 cells, where PFD reduced doxorubicin resistance (Fig. S5H).

TNFα activates the NFκB pathway that increases anti-apoptotic gene expression to promote cell survival (57–59). Our observation of early activation of p38 MAPK-MK2 (Fig. 4A-B) suggested that TNFα could be rapidly up-regulated upon G0 induction. Time-course translatome analysis affirmed that TNFα is highly increased (16-fold) at the earliest time point of 4 h after serum-starvation or AraC treatment (Fig. 5B) along with its receptors, leading to rapid elevation of downstream NFκB target genes including anti-apoptotic BCL family members (59–61) (Fig. 5B, S5A-B). Similar to our observations with TNFα inhibitor PFD, NFκB inhibitor, BAY11-7082(62) prior to or along with AraC or SS decreases survival of G0 cells, while treatment after AraC or SS had no effect (Fig. 5H). These data suggest that the TNFα-NFκB inflammatory pathway is upregulated as an early survival pathway in G0 cells.

### TTP regulates a pro-apoptotic JNK pathway via targeting DUSP1

We asked what other ARE mRNAs are targeted by TTP and affect cell survival. DUSP1 mRNA contains AREs in its 3’ UTR. TTP has been shown to target DUSP1 mRNA for degradation upon LPS treatment of macrophages or dendritic cells (44;45;63). To determine if TTP phosphorylation regulates DUSP1 in AraCS, we expressed TTP-AA mutant that is not phosphorylated in BMDM cells that lack TTP (Fig. 6A). Immunoprecipitation showed that TTP-AA associated with DUSP1 mRNA in AraCS cells (Fig. 6B). Expression of TTP-AA mutant more significantly reduced DUSP1 mRNA and protein levels compared to cells expressing TTP wild-type (Fig. 6C-D). Furthermore, inhibition of phosphorylation of TTP by p38 MAPK inhibitor decreased DUSP1 protein level (Fig. 6E). DUSP1 is a MAPK phosphatase which dephosphorylates JNK (64). In AraCS cells, DUSP1 protein level is negatively correlated with phosphorylated JNK (Fig. 6C, 6E), consistent with DUSP1-mediated suppression of JNK(64). To determine the effect of JNK on survival of leukemic cells, JNK inhibitor, JNK-IN-8 was used (Fig. 6A). Importantly, JNK inhibition reversed apoptosis of leukemic cells treated with AraC, LY and PFD, but did not affect the viability of untreated cells (Fig. 6F), indicating that inhibition of JNK pathway contributes to chemoresistance. Together, these results suggest that TTP-DUSP1 axis promotes chemoresistance via suppressing JNK-mediated apoptosis (Fig. 6A).

**Figure 6.**
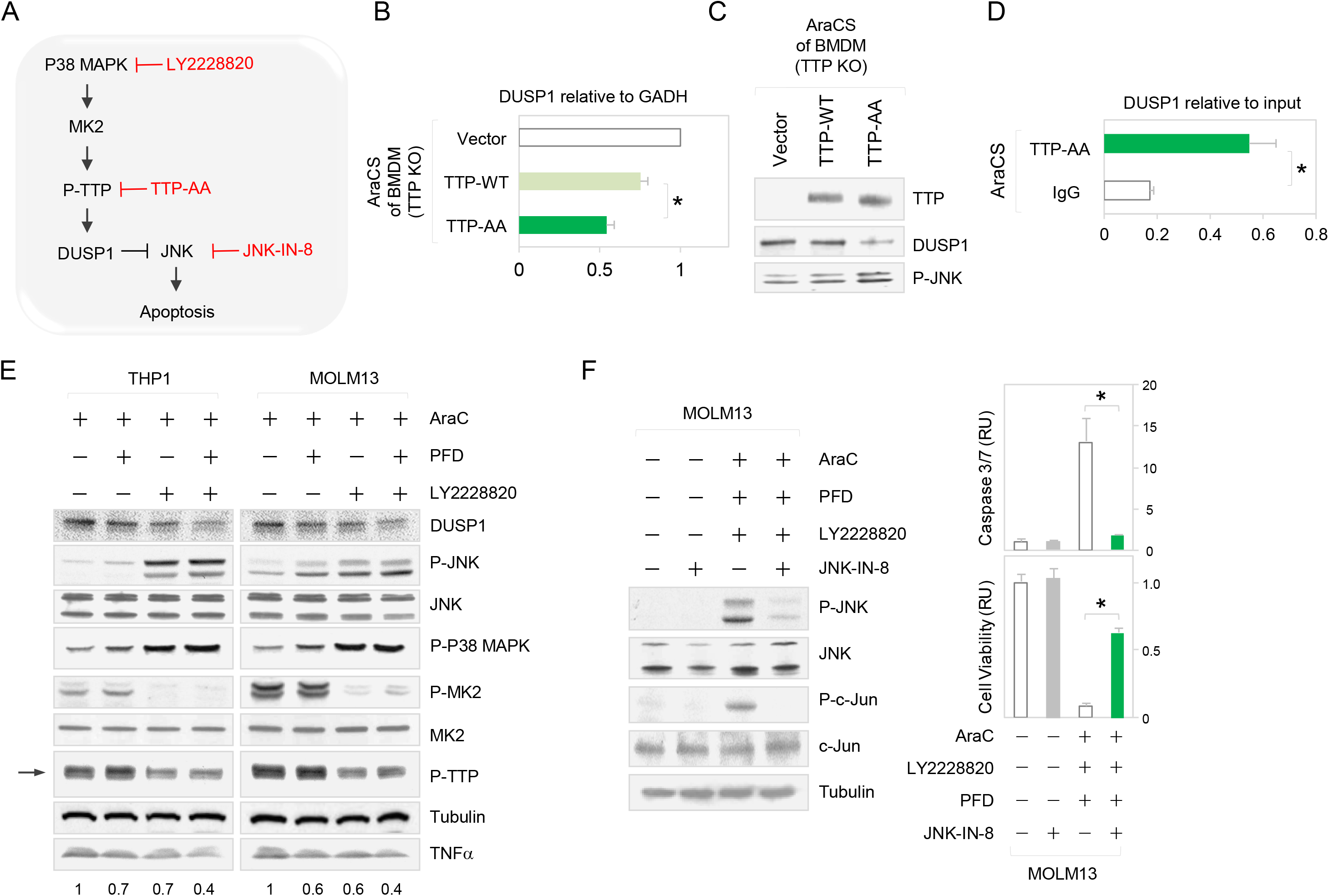
TTP regulates a pro-apoptotic JNK pathway via targeting DUSP1. **A**. Phosphorylation of TTP allows expression of the ARE-bearing mRNA of DUSP1 that inhibits JNK, and hence blocks JNK-mediated apoptosis. JNK pathway is blocked by the inhibitor JNK-IN-8. **B-D**. Effect of TTP-AA mutant on DUSP1 and phosphorylation of JNK. BMDM TTP deficient cells were treated with doxycycline to express TTP-AA and TTP wild-type prior to AraC treatment. DUSP1 mRNA level was measured by qPCR and is shown relative to GAPDH mRNA (B). Western analysis of TTP, DUSP1 and phospho-JNK is shown (C). TTP-AA was immunoprecipitated with GFP antibody, followed by qPCR analysis for DUSP1 mRNA (D). **E**. Western analyses in THP1 and MOLM13 cells treated with indicated drug combinations. Phospho-TTP is indicated with an arrow and quantitation of TNFα protein is shown below. **F**. JNK pathway mediates apoptosis. MOLM13 cells treated with indicated drug combinations. JNK pathway was inhibited with 1 μM JNK-IN-8. Western analyses of phospho-JNK, phospho-c-Jun and c-Jun shown on the left; associated cell viability and death graphed on the right. Data are represented as average ± SEM.

### Co-inhibition of p38 MAPK and TNFα sensitizes resistant leukemic cells to AraC treatment

Although chemoresistant cells are sensitive to individual inhibition of either TNFα or p38 MAPK by PFD or LY respectively, a substantial proportion of cells still survived (Fig. 4F, 5G). Therefore, we asked if coinhibition of p38 MAPK and TNFα with LY *and* PFD respectively, could eliminate the remaining resistant cells. We find that individual treatment with either of LY or PFD prior to or along with AraC, reduces approximately 50% of surviving leukemic cells (Fig. 7B). Importantly, the combination of <u>P</u>FD *and* <u>L</u>Y2228820 prior to <u>A</u>raC treatment–called PLA therapy–eliminates about 90% of chemoresistant cells in multiple AML cell lines (Fig. 7A-C). Furthermore, PLA therapy decreased colony formation capacity of leukemic cells on methylcellulose by 10-fold, compared to AraC-treatment alone (Fig. 7D). These data indicate a severe loss of stem cell capacity of leukemic cells treated with PLA therapy. In contrast, in the absence of AraC treatment, the combination of PFD and LY2228820 did not affect cell viability, apoptosis and colony formation capacity, indicating the synergistic effect between AraC and anti-inflammatory drugs (Fig. 7B-D). Despite the fact that stromal niche cells have been shown to protect leukemic cells from chemotherapy (65), we find that AML cells co-cultured with stromal cells remained sensitive to PLA therapy (Fig. S5E). We investigated the molecular mechanism by which PLA therapy enhanced chemosensitivity. We find that LY treatment destabilizes TNFα mRNAs by TTP dephosphorylation (43) (Fig. 3G, 4C), while PFD suppresses translation of TNFα mRNA (56) (Fig. 5E). Therefore, in PLA therapy, TNFα remains more effectively blocked, compared to individual drug treatments (Fig. 6E). Furthermore, a pro-apoptotic JNK pathway was more significantly activated in cells treated with PLA therapy than single-drug treatments (Fig. 6E). Together, these results suggest that PLA therapy reduces TNFα and promotes a pro-apoptotic JNK pathway, leading to apoptosis of chemoresistant cells.

**Figure 7.**
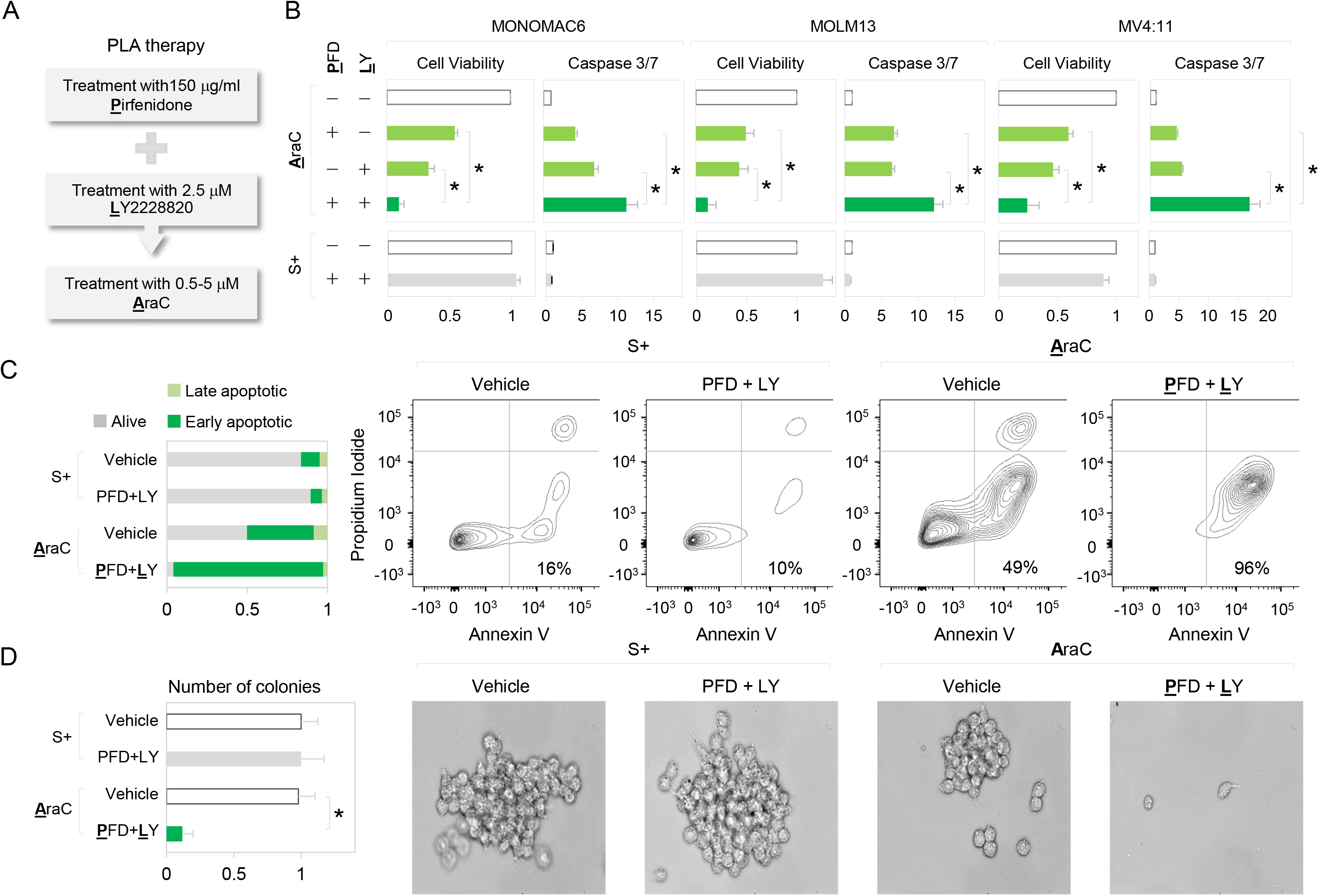
PLA therapy decreases AraC-resistant cells in AML cell lines. **A**. PLA therapy, involves pre-treatment of leukemic cells with <u>P</u>FD and <u>L</u>Y followed by <u>A</u>raC treatment, using half of the concentrations used for individual drugs in Fig. 4 and 5. **B**. Three different AML cell lines were sequentially treated with indicated drugs, followed by assessment of cell viability and death. **C-D**. Viability of MOLM13 cells treated with indicated drug combinations. Flow cytometric profiles of cells stained with annexin V and propidium iodide are shown (C). Cells were plated on methylcellulose media to test colony formation in the presence of drug combinations. Representative colony images and quantification of colonies are shown (D). \**P*□≤□0.05. Data are represented as average ± SEM. See also Fig. S5.

### PLA therapy reduces chemoresistance in primary AML cells *ex vivo* and *in vivo*

To test the anti-leukemic activity of PLA therapy in primary AML (66), primary cells from AML patients as well as two murine AML models driven by Hoxa9/Meis1 or MLL/AF9 (SI Methods), were used. When either p38 MAPK or TNFα was inhibited prior to AraC treatment, moderate apoptosis of chemoresistant cells was observed in primary AML cells (Fig. 8A-B). Importantly, co-inhibition of p38 MAPK and TNFα by PLA therapy (pre-treatment before AraC) significantly reduced AraC resistance in fourteen out of fifteen AML patient samples as well as in primary cells from two AML mouse models *ex vivo* (Fig. 8A-B). In contrast, the viability of normal CD34+ cells from healthy donors was not affected by treatment with LY or PFD (Fig. 4J, 8A-B), consistent with clinical studies that have shown that PFD and LY have acceptable safety and tolerance (53;55). To further investigate the therapeutic potential of PLA therapy *in vivo*, human AML cells expressing luciferase (MOLM13-Luc, SI Methods) were intravenously or subcutaneously injected into NSG mice. After confirmation of engraftment by measuring tumor volume or bioluminescent imaging (BLI), the mice were treated with PLA therapy or AraC for two weeks. Consistent with *ex vivo* results (Fig. 7B), PLA therapy significantly decreased the leukemic burden and tumor volume by 6-fold, compared to AraC treatment alone (Fig. 8C-D). Next, primary Hoxa9/Meis1 or MLL/AF9 leukemia cells were generated as described previously (67), and transplanted to second recipient mice. These mice were treated with PLA therapy or AraC. Consistently, BLI shows that PLA therapy eliminated 78% or 96% of chemoresistant cells in a dosage-dependent manner (Fig. 8E-F) and extended mice survival (Fig. 8H and S5G). In the absence of AraC treatment, the combination of PFD and LY2228820 did not affect leukemic burden, suggesting that cytotoxic effects of this combination are limited to AraC-resistant cells, rather than proliferating cells (Fig. 8G). Together, these results suggest PLA therapy has potential for improving AraC-mediated apoptosis in AML.

**Figure 8.**
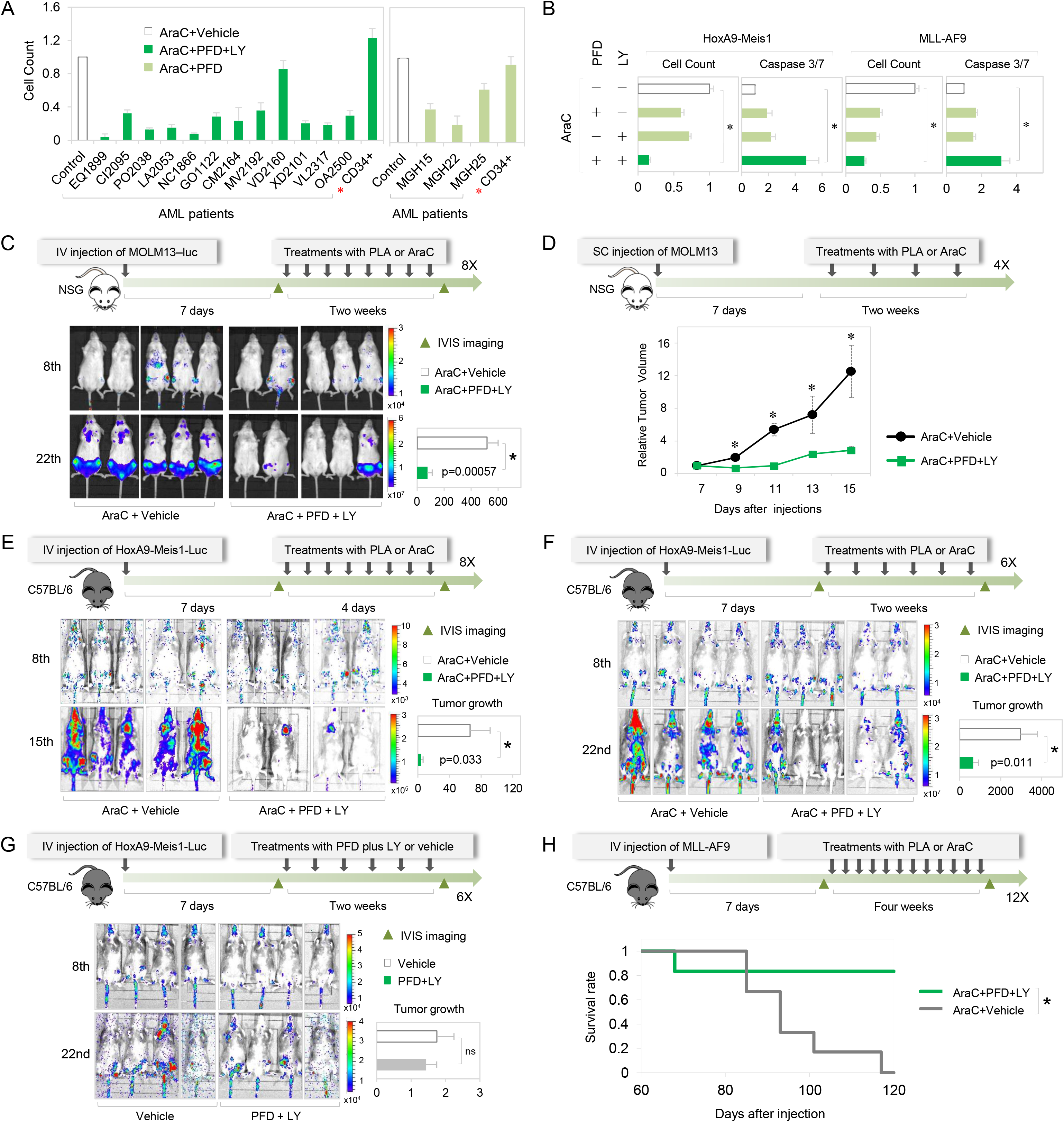
PLA therapy significantly reduces AraC resistance in primary AML cells *ex vivo* and *in vivo*. **A**. Viability of primary cells from AML patients and normal CD34+ cells from healthy donors after indicated treatments. **B**. Viability and death of primary cells from AML mouse models driven by HoxA9-Meis1 and MLL-AF9 after indicated treatments. **C-G**. Bioluminescence images and quantification of tumor growth in NSG mice engrafted with MOLM13 cells and treated with PLA therapy or AraC (C-D) and in C57BL/6 mice engrafted with primary HoxA9-Meis1/luciferase cells and treated with PLA therapy or AraC (E-F) or treated with PFD plus LY or vehicle as a control (G). **H**. Kaplan-Meier survival curves of MLL-AF9 engrafted C57BL/6 mice, treated with PLA therapy or AraC. \**P*□≤□0.05. Data are represented as average ± SEM. See also Fig. S5-S6.

## Discussion

G0 cells are a transiently arrested, clinically relevant subpopulation in cancers (1;2;5–10). Our previous data and others, revealed altered gene expression mechanisms in G0 leukemic cells, at the post-transcriptional (8;12) and translational levels (13;14;18). This would lead to a distinct gene expression profile to enable G0 cell survival in harsh conditions. G0 cells are resistant to stress conditions like serum-starvation, with transient inhibition of apoptosis, and proliferation (1;11;18). Importantly, we find that serum-starved leukemic SS G0 cells exhibit chemoresistance (Fig. 1E); consistently, true chemo-surviving AraCS cells are transiently arrested and chemoresistant (Fig. 1D-E, S1B-C). In accord, we find that SS cells are similar in translatome and proteome to AraCS cells (Fig. 1F), indicating that consistent with their common features of G0 arrest and chemosurvival, they show similar post-transcription gene expression. Published transcriptional signatures of in vivo chemoresistance leukemic models (1;2;8;11;15;16), are also highly expressed in SS and AraCS cells (Fig. 1G, S1G). Thus, the common G0 resistance gene expression profile observed in AraCS and SS G0 cells likely comprises genes that control survival and resistance. These data revealed that in addition to known transcriptional profiles, altered post-transcriptional mechanisms in G0 resistant cells contribute to their unique gene expression profile that underlies their chemoresistance.

Our findings reveal the importance of DNA damage and stress signaling that can initiate a pro-inflammatory response that causes survival (Fig. 4). Differential genomic instability in cancers would lead to subpopulations within a tumor with disparate DDR and stress signaling (47–49) that we find, enables their chemotherapy survival via pro-inflammatory cytokines. Cytokines upregulated in SS and AraCS cells include some SASP factors but also other unique cytokines (36;37) (Fig. S2G). This is consistent with similarities and differences between G0 and senescence (1): both show low mTOR activity but G0 shows reversible arrest, stem cell markers, low p53 and lack of senescence markers (Fig. 2C-D, 2I, S1G) (14)—unlike senescence(18). These data indicate that a quiescence- and resistance-specific set of pro-inflammatory and signaling genes are expressed in these resistant cells (Fig. 2G). These include inflammatory cytokine, TNFα that promotes downstream NFκB activated prosurvival target genes (57–59) including BCL family members of anti-apoptotic genes (59–61) (Fig. 5A-C, S5A-B). Treatment with anti-inflammatory reagents *after* chemotherapy is not very effective as the downstream survival effectors have already been induced; thus, targeting their upstream cytokine regulators would not be effective at this later time (Fig. 4F-G, 5F-H, S5D). Therefore, treatment with reagents that block these resistance pathways *prior to* (and continued with) or along with chemotherapy, enables the most effective reduction of resistance, as they prevent further enrichment of such resistant cells by blocking induction of pro-survival signaling.

Increasing AraC, a nucleotide analog that inhibits replication (17), would activate DDR and downstream p38 MAPK signaling (47–49) and should lead to more cells expressing this inflammatory pathway that enables resistance. Consistently, increased AraC treatment leads to more cells in the inflammatory phase that can be targeted by LY to curb resistance (Fig. 4K). Non-cancerous cells do not show this pathway (Fig. 2I) and are not affected by inhibitors (Fig. 4J, 8A). These data suggest that certain chemotherapies and stresses like serum-starvation induce stress signaling (Fig. 4A-C) and enrich for resistant G0 cells—in addition to pre-existing subpopulations with genomic instability that trigger DDR and stress (47–49). Importantly, this resistance mechanism can be blocked, not only in different AML cell lines (Fig. 4J, 5G, 7B) but also in vivo (Fig. 8C-G) and in multiple patient-derived primary AML—without affecting normal cells (Fig. 8A)—supporting their potential applicability as a therapeutic against chemoresistance in AML.

We find key signaling pathways induced by AraCS and SS treatments, which alter post-transcriptional and translational gene expression to enable resistance. These include: 1. DNA damage ATM (47–49) and stress activated p38 MAPK signaling that in turn promotes MK2 (51;52) that post-transcriptionally upregulates ARE bearing mRNAs (43;45;46). The expressed mRNAs include ARE-bearing proinflammatory cytokine TNFα (57;58) that activates downstream anti-apoptosis signals (Fig. 4A-D, 5A-C, S5A-B) (59–61), and ARE-bearing signaling regulator DUSP1(63;64)that blocks JNK-mediated apoptosis (Fig. 6), to promote resistance. 2. ATM-mediated suppression of mTOR activity (47;48) that inhibits canonical translation initiation via 4EBP dephosphorylation (Fig. 2A-C); this results in specific translation of pro-inflammatory cytokines(14) (Fig. 3A-C) and immune modulators (30) (HLA-G, CD47, Fig. 2F, S2I) (31–33) that regulate anti-tumor immune response and resistance (34;35). 3. UPR stress signaling, induced downstream of p38 MAPK (68) and DNA damage (69;70), also inhibits canonical translation via PERK phosphorylation of eIF2α and enables non-canonical specific mRNA translation (Fig. 2A-D, S2H)(69;70). Blocking the p38 MAPKα/β pathway with LY (52;53), in combination with the anti-inflammatory PFD (52;55;56) that precludes downstream TNFα expression (55;56) (Fig. 5E)—prior to (and continued with) AraC chemotherapy—lead to effective loss of chemoresistance in multiple AML cell lines (Fig. 7B), in tumors *in vivo* in AML mouse models (Fig. 8C-G), and in patient samples (Fig. 8A), validating their ability to reduce resistance and tumors *in vitro* and *in vivo*. LY destabilizes TNFα mRNA by TTP dephosphorylation (Fig. 4C) (43), while PFD suppresses TNFα selectively at the translation level (56) (Fig. S5F) and thus enables PLA combination therapy to more effectively curb resistance than the individual drugs (Fig. 7B, 8B). Apart from its effect on TNFα translation, PFD blocks inflammation regulator (71;72) p38 MAPKγ that can be increased upon p38MAPKα/β inhibition, preventing feedback reactivation of inflammation, and enabling PLA combination therapy to remain more efficacious than the individual drugs. Therefore, the combination of PFD and LY suppresses the inflammatory and stress response more effectively *in vitro* and *in vivo* (Fig. 7-8). Upon inhibition of p38 MAPK, in addition to reduction of TNFα and its downstream anti-apoptotic signals, we find the ARE bearing DUSP1 is reduced, leading to activation(63;64)of the JNK pathway(73) to promote apoptosis (Fig. 6E-F). These data indicate that blocking pro-inflammatory effectors—that are induced by chemotherapy mediated DNA damage and stress signaling—leads to increased chemosensitivity and decreased resistant cell survival.

Our findings revealed that these pro-inflammatory and signaling genes upregulated in G0, have AREs and other UTR sequences that regulate mRNA levels and translation (Fig. 3A-C, S3A). The ATM-p38 MAPK-MK2 axis stabilizes these ARE bearing pro-inflammatory cytokine and signaling mRNAs by phosphorylating ARE binding mRNA decay factor, TTP to prevent its mRNA decay activity on pro-inflammatory cytokine TNFα (Fig. 3D-H, 4C-D) and signaling regulator, DUSP1 (Fig. 6A-D). In support, overexpression of TTP-AA—that cannot be phosphorylated and is a dominant active form that restores ARE mRNA decay (43–45)—decreases TNFα and DUSP1 expression (Fig. 3F-G, 6A-D), and thereby reduces chemoresistance (Fig. 4H-I, 6E-F). This is consistent with previous studies on AREs in cancers (14;38;43;74–77). These data suggest that phospho-TTP level or TTP activity is an important regulator of inflammatory response mediated chemoresistance, which can be harnessed as a marker and target against AML resistance. Consistently, published in vivo leukemia resistance models show increased expression of TTP and ARE bearing genes (15;78), similar to our studies (Fig. 3A-E). Our studies on TTP and ARE regulated immune and signaling modulators that promote chemoresistance, are consistent with recent findings of TTP regulation of PDL1 to mediate immunoresistance in solid tumors(79). Importantly, inhibition of these pathways curtails chemoresistance and tumor survival *in vivo* in primary AML patients and tumor models (Fig. 8). Together, these pathways that are upregulated in resistant cells (Fig. 4A, 5A) via chemotherapy and stress induced signaling—decrease canonical translation and permits non-canonical post-transcriptional regulation of specific genes (Fig. S6)—to promote chemosurvival of G0 cancer cells.

## Conclusions

Our studies reveal that G0 leukemic cells are chemoresistant, indicating their clinical importance in cancer persistence. We find a specific proteomic and translation profile that is induced commonly between G0 cells and chemosurviving leukemic cells. We uncovered critical genes that are specifically upregulated post-transcriptionally and translationally for cell survival in these conditions by key survival signaling pathways. These studies reveal the significance of post-transcriptional and translational regulation of immune and signaling modulators in chemoresistance. Our data enabled the development of a new combination therapy to effectively reduce resistance in cancer cell lines, in tumors in vivo, and in patient tumor samples, without affecting normal cells.

## Methods

### Overview, aim, design, and setting

Therapeutic targeting of minimal residual disease or chemoresistant, leukemic stem cells in leukemias, particularly acute myeloid leukemia, has been ineffective thus far and refractory leukemia is fatal. The mechanisms of translation and post-transcriptional control, and the critical translation profile that control the ultimate, specific protein profile, and thereby—survival of such clinically resistant cells—are largely undiscovered. Therefore, we globally analyzed gene expression at every level—RNA levels, translatome and proteome—in chemotherapy-surviving G0 cancer cells in acute monocytic leukemia and other cancers, the specialized post-transcriptional and translational mechanistic changes, their key signaling regulatory pathways, as well as developed a new, resistance-gene expression targeting therapy to understand and reduce chemoresistance.

**Detailed description of characteristics, materials used, and methods** including cell culture, patient samples, tumor models, profiling, plasmids, cell viability assays, flow cytometry, protein analysis, drugs, and motif analysis are described in detail in Supplemental Information.

**Statistical analyses** are described in Supplemental Information.

## Declarations

### 1. Ethics approval and consent to participate

**Statement on Human Data** All human samples (de-identified) were handled in accordance with IRB protocols to SV (2015P000998/MGH), approved by the Partners Human Research Committee Institutional Review Board /MGH IRB, to DAS, and to Tim Graubert & J. L-S (DF/HCC 13-583), approved by DF/HCC Office for Human Research Studies. Details in Supplemental Information.

**Statement on animal data** nod-scid-gamma (NSG), C57Black/6 mice, 10-12 weeks, are obtained from MGH Cox-7 Gnotobiotic animal facility of the AAALAC-accredited Center for Comparative Medicine and Services at MGH. These facilities are supervised by veterinarians in the Center for Comparative Medicine and the MGH Subcommittee for Animal Research (SRAC) and maintained according to the protocol approved by SRAC, and provide services for breeding, regular health checks, histopathology and macropathology.

### 2. Consent for publication

All samples are de-identified, described under the above IRB protocols.

### 3. Availability of material and data

Raw datasets will be submitted to GEO public repository at final submission. All datasets and material will be made available publicly on publication and on request.

### 6. Author Contributions

SL conducted the research and bioinformatic analysis; SST, SIAB & DL contributed data; SL, YK, DM, BTN, ID-G, DTM, M-KC, DS, SM & DAH provided in vivo models & in vivo data; MAM, RR & RG did immune data; CT & MD-M provided TTP reagents & stable knockdown cells, AL, NJH, & ML provided mRNA folding energies & patient gene signatures; MB & WH conducted proteomics; DAS & JL-S provided patient samples; SV supervised the project & wrote the manuscript.

## 7. Acknowledgments

We thank Partners Healthcare Center for Personalized Genetic Medicine & BUMC facilities for microarray data; N. Kedersha, S. Lyons & P. Anderson for plasmids & antibody; T. Graubert for patient samples; D. Bloch, S. Wu, A. Naar, M. Gaestel, M. Guzman, N. Bardeesy, D. Scadden, S. Ramaswamy, & C. Benes for reagents.

## List of Supplemental files

1. **Supplemental Information** Methods (related to main text, main figures 1-8 & supplemental figures S1-S6) Supplemental References (related to supplemental information, methods section) Supplemental Figure legends S1-S6, related to main figures 1-8
2. **Supplemental Figures S1-S6 (related to main figures 1-8)**
3. **Supplemental Tables 1-2 (related to main figures 1-8 & supplemental Figures S1-S6)**

