## Supplementary figures and images for "A post-transcriptional program of chemoresistance by AU-rich elements and TTP"

### SI_Figures

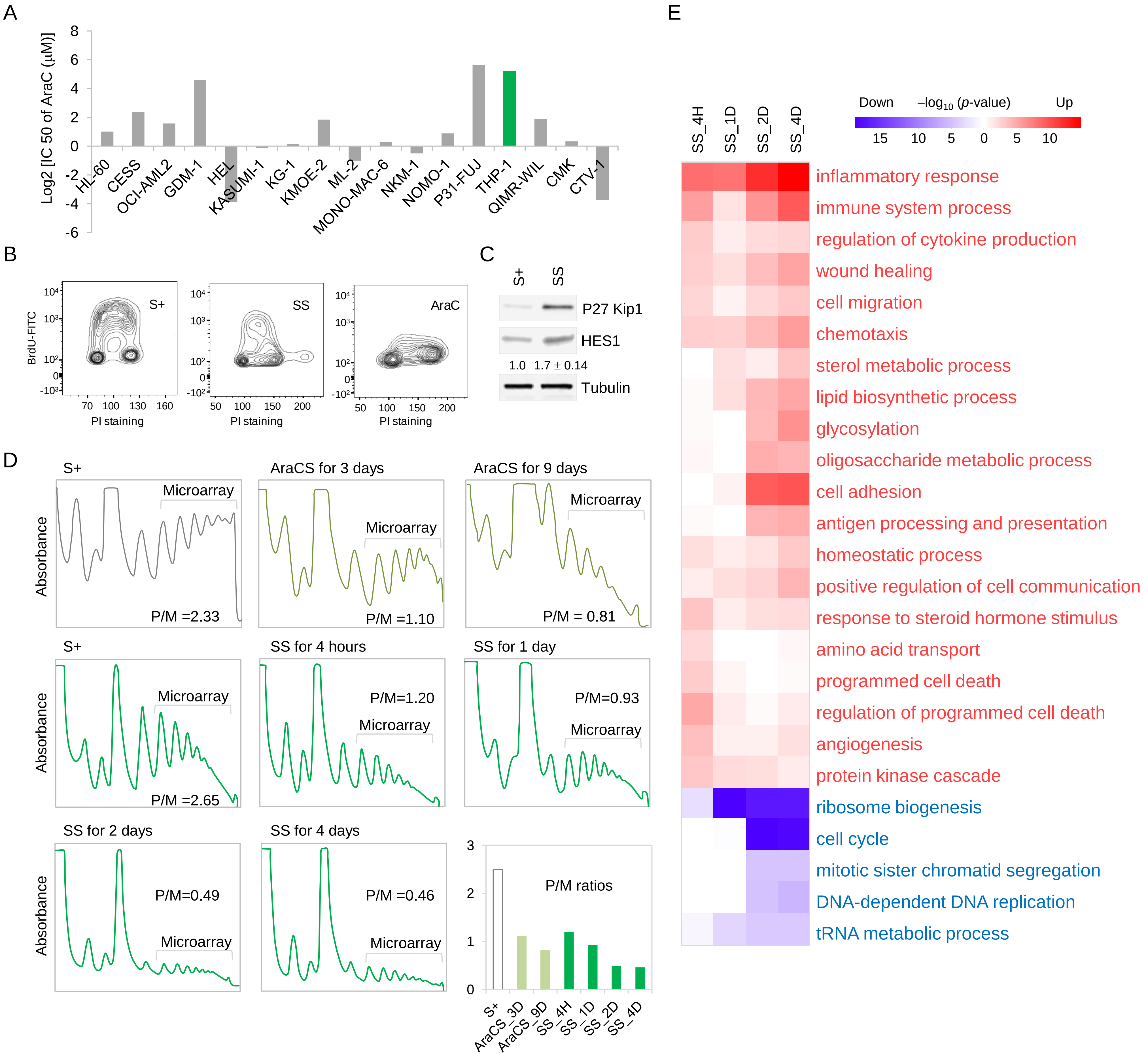

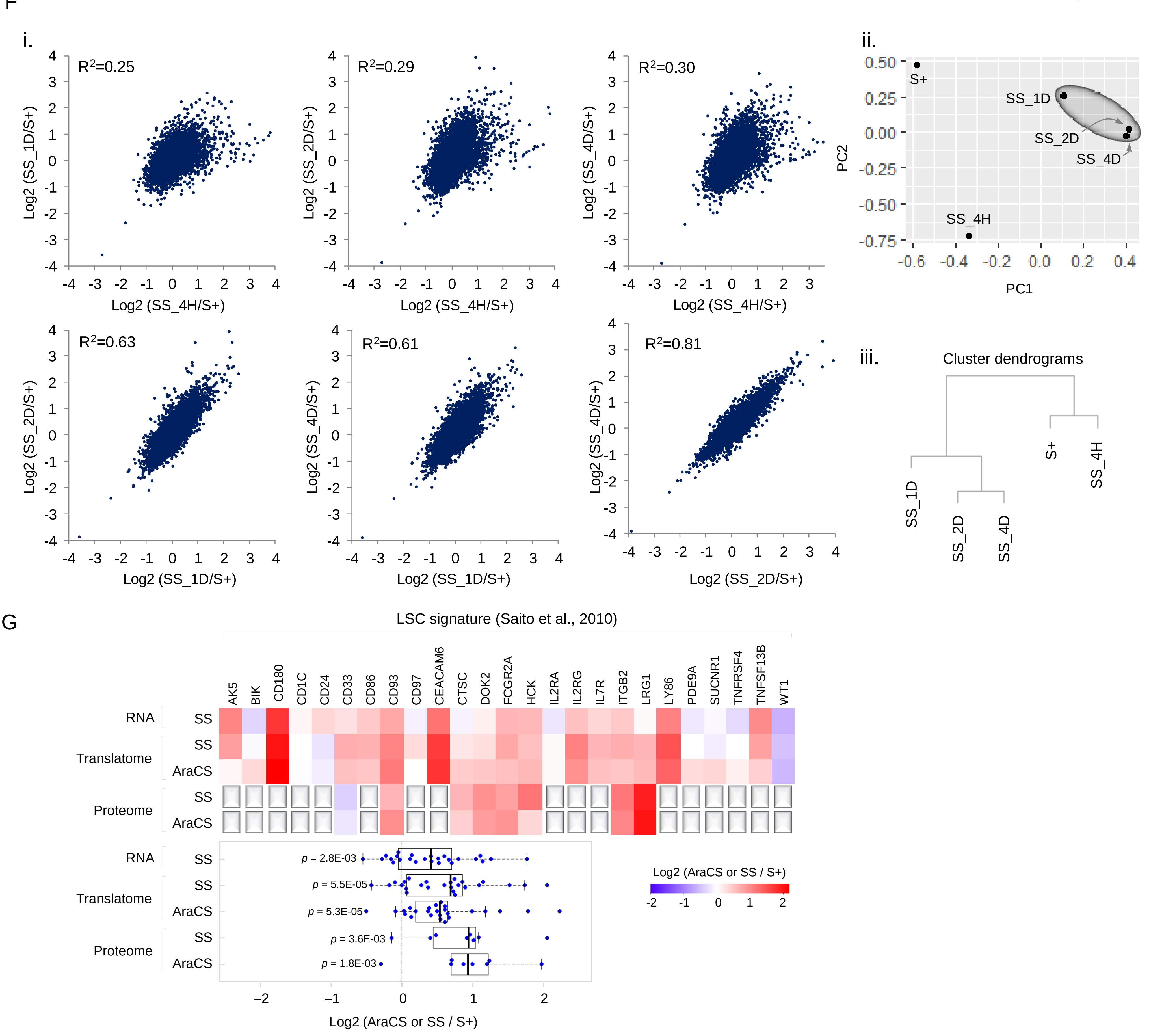

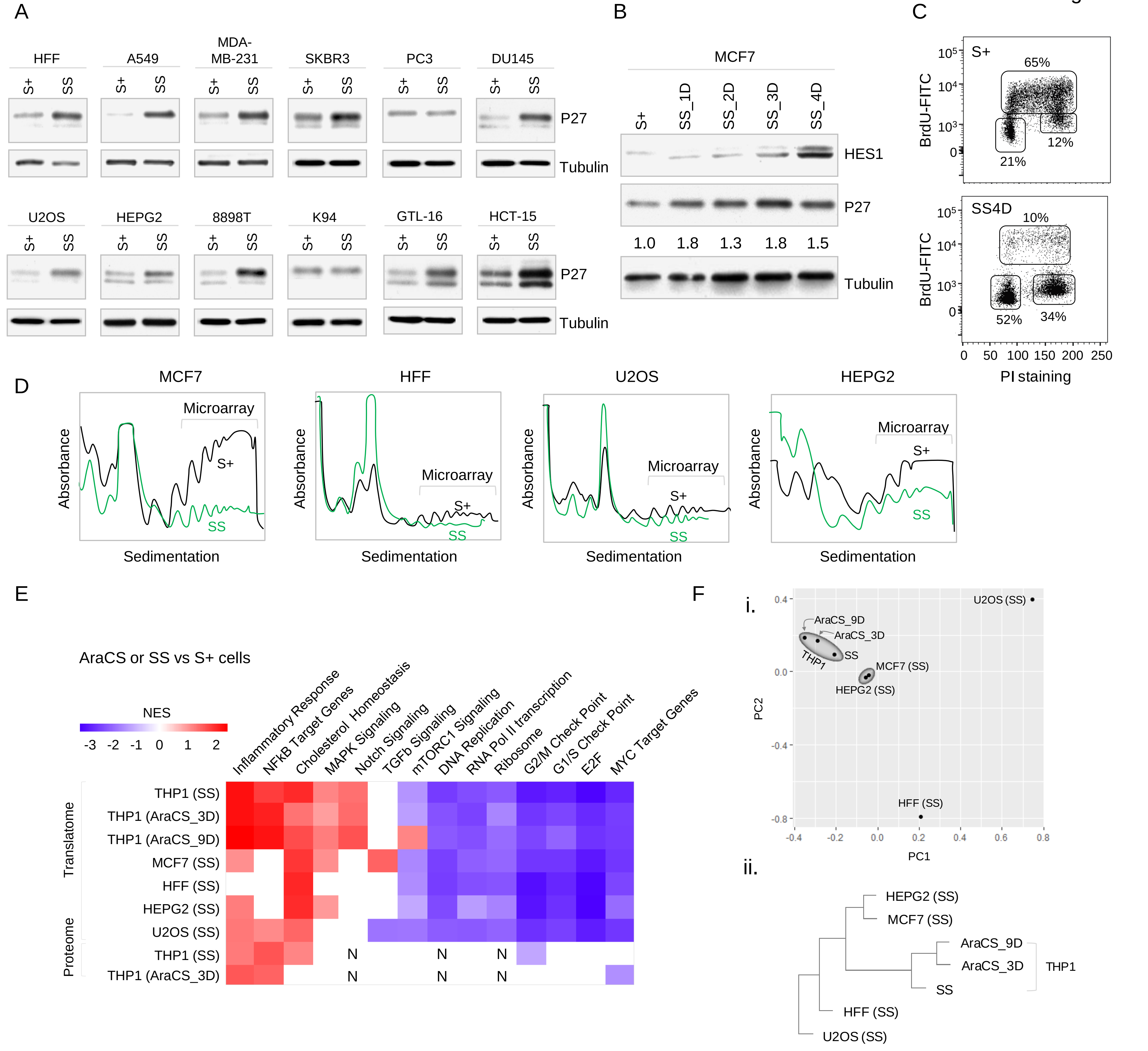

Figure S2

G

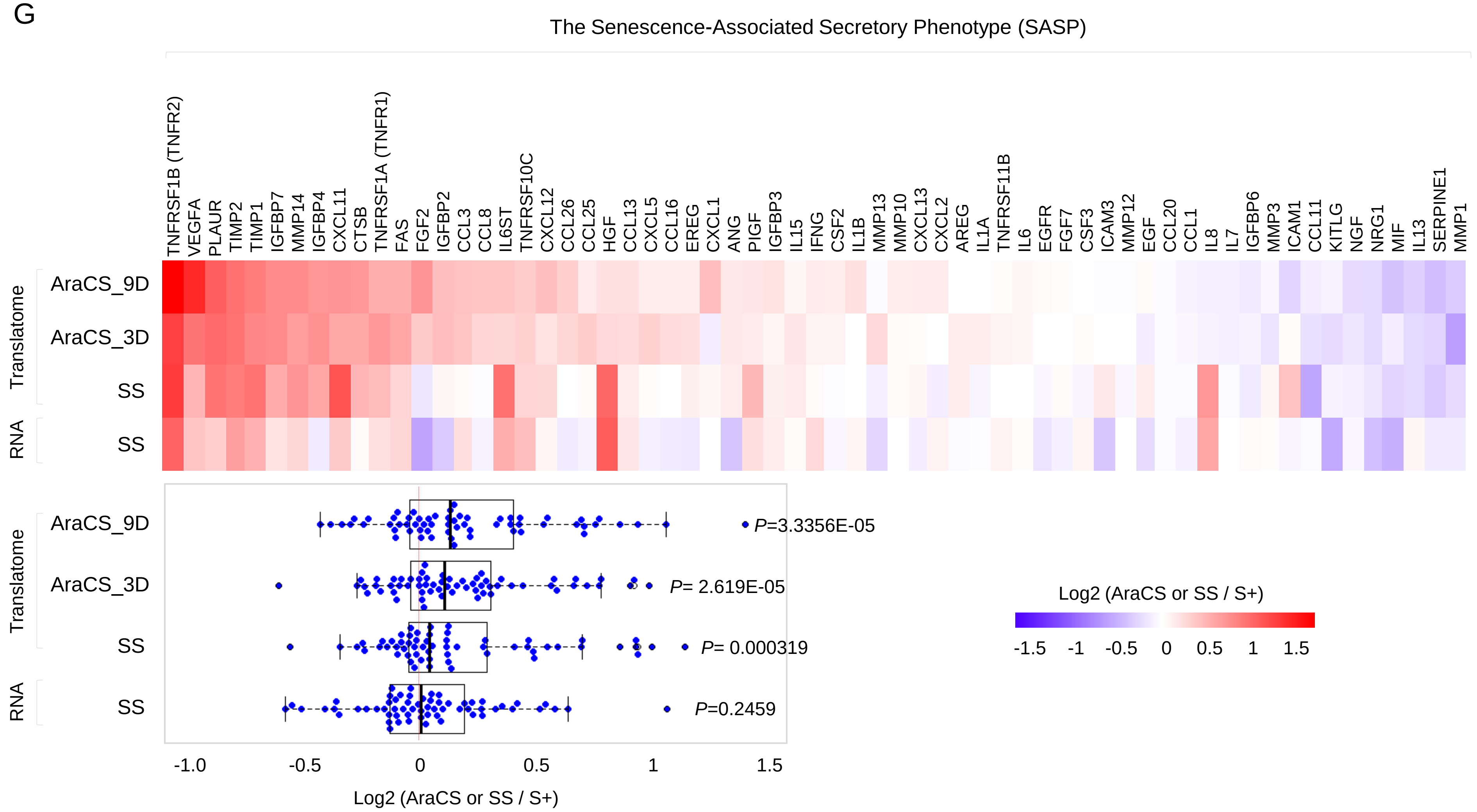


H

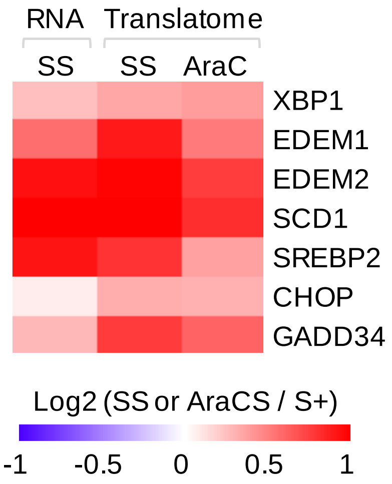


I

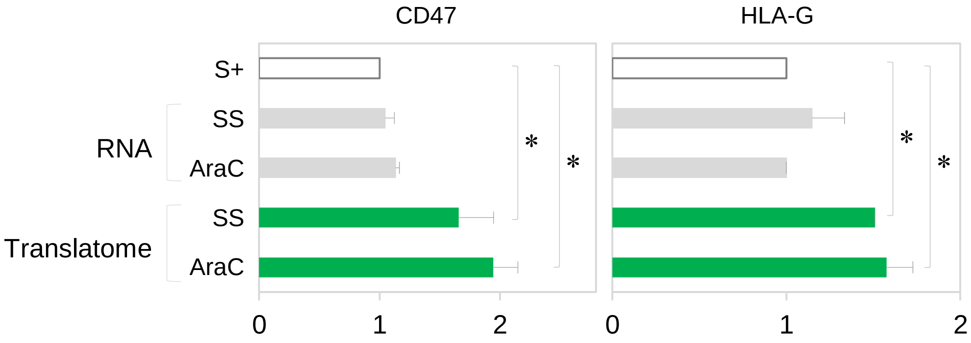

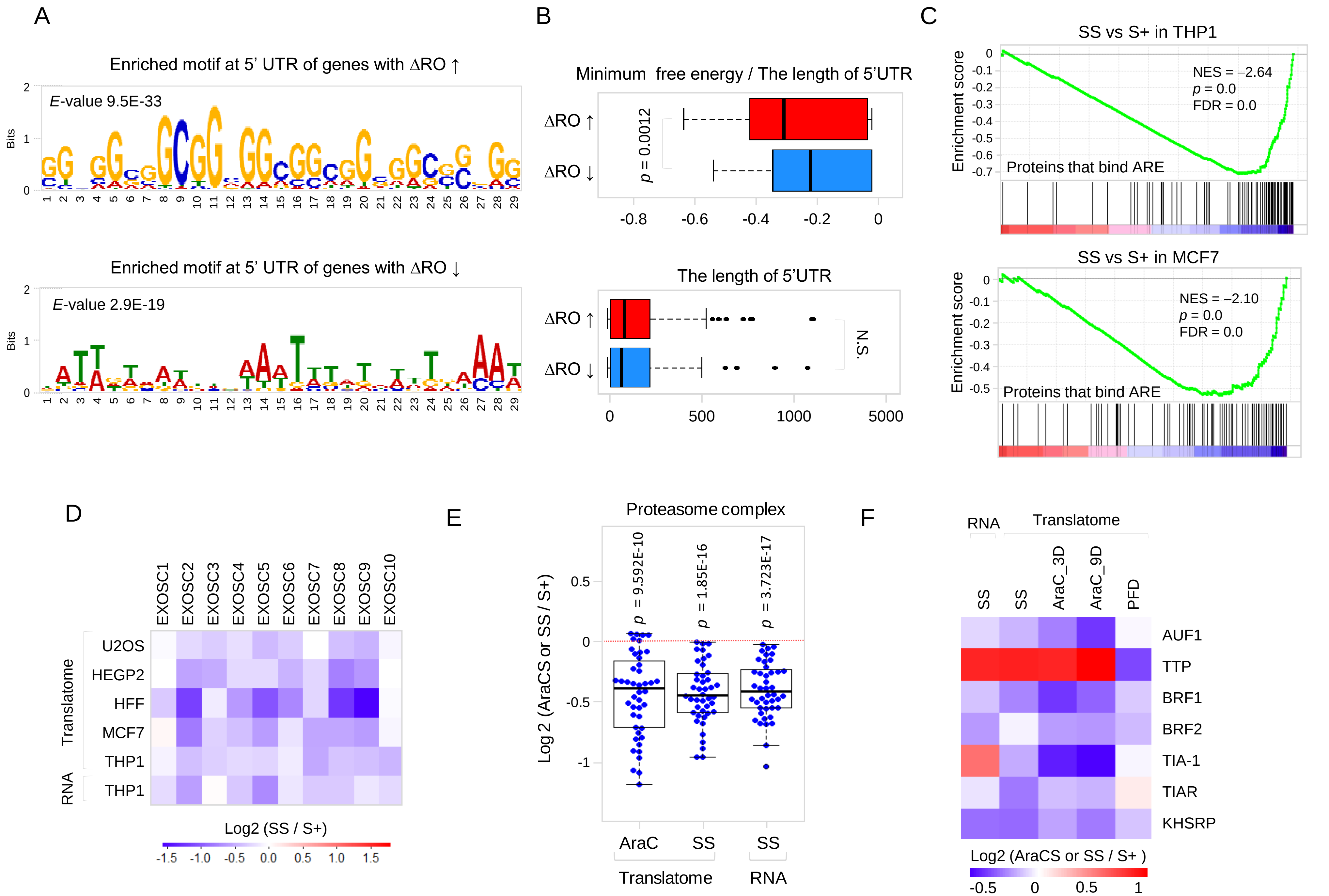

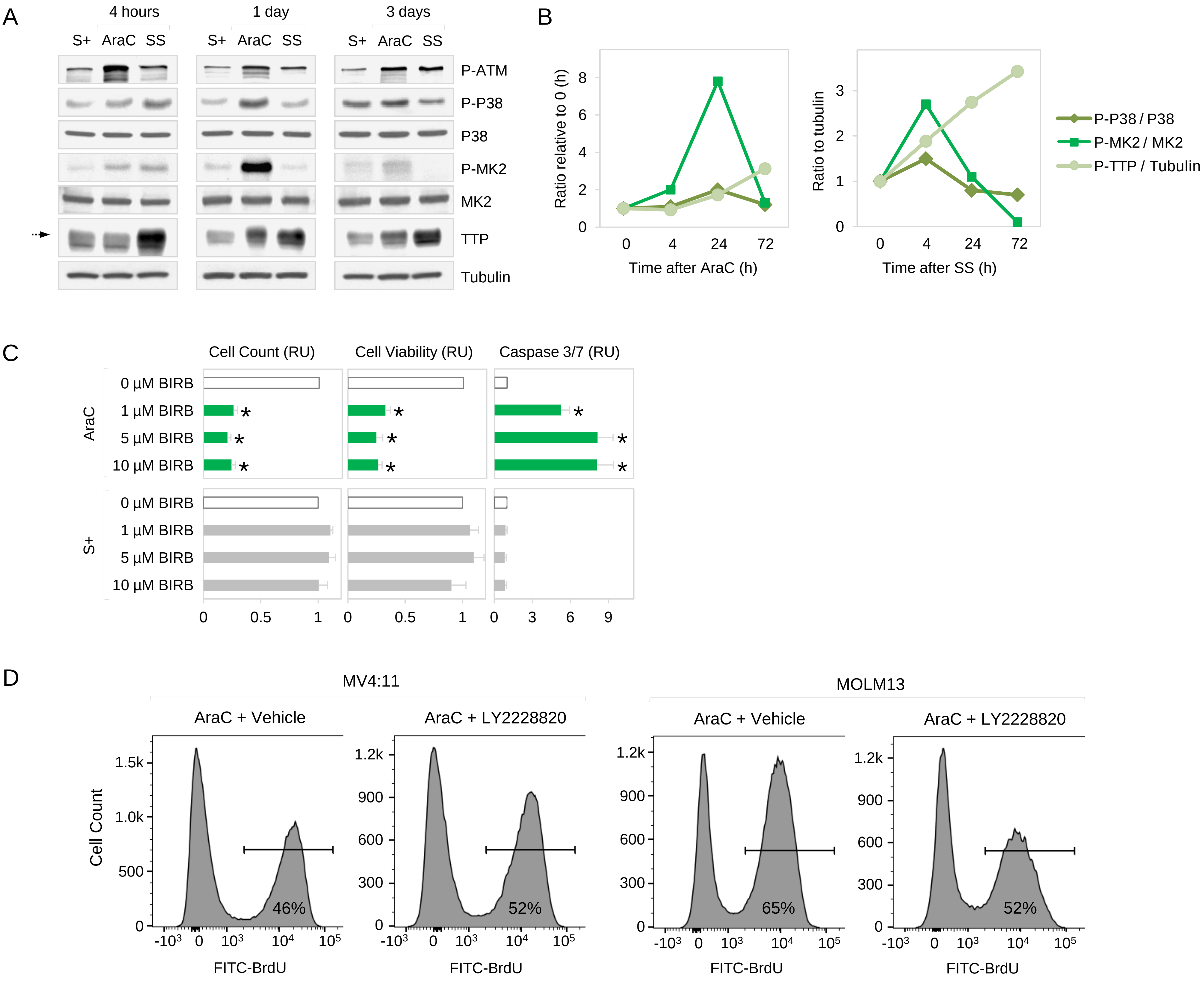

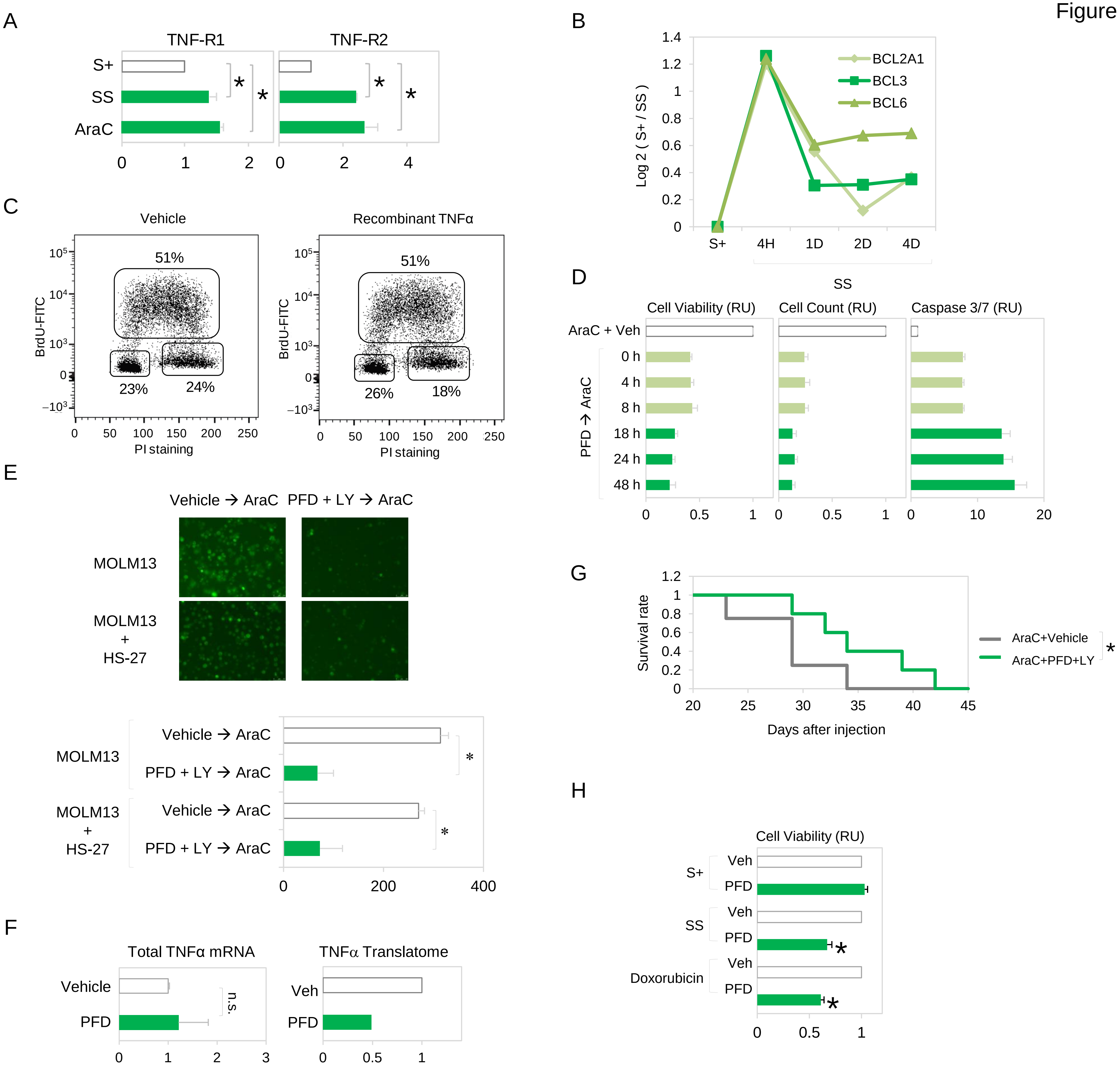

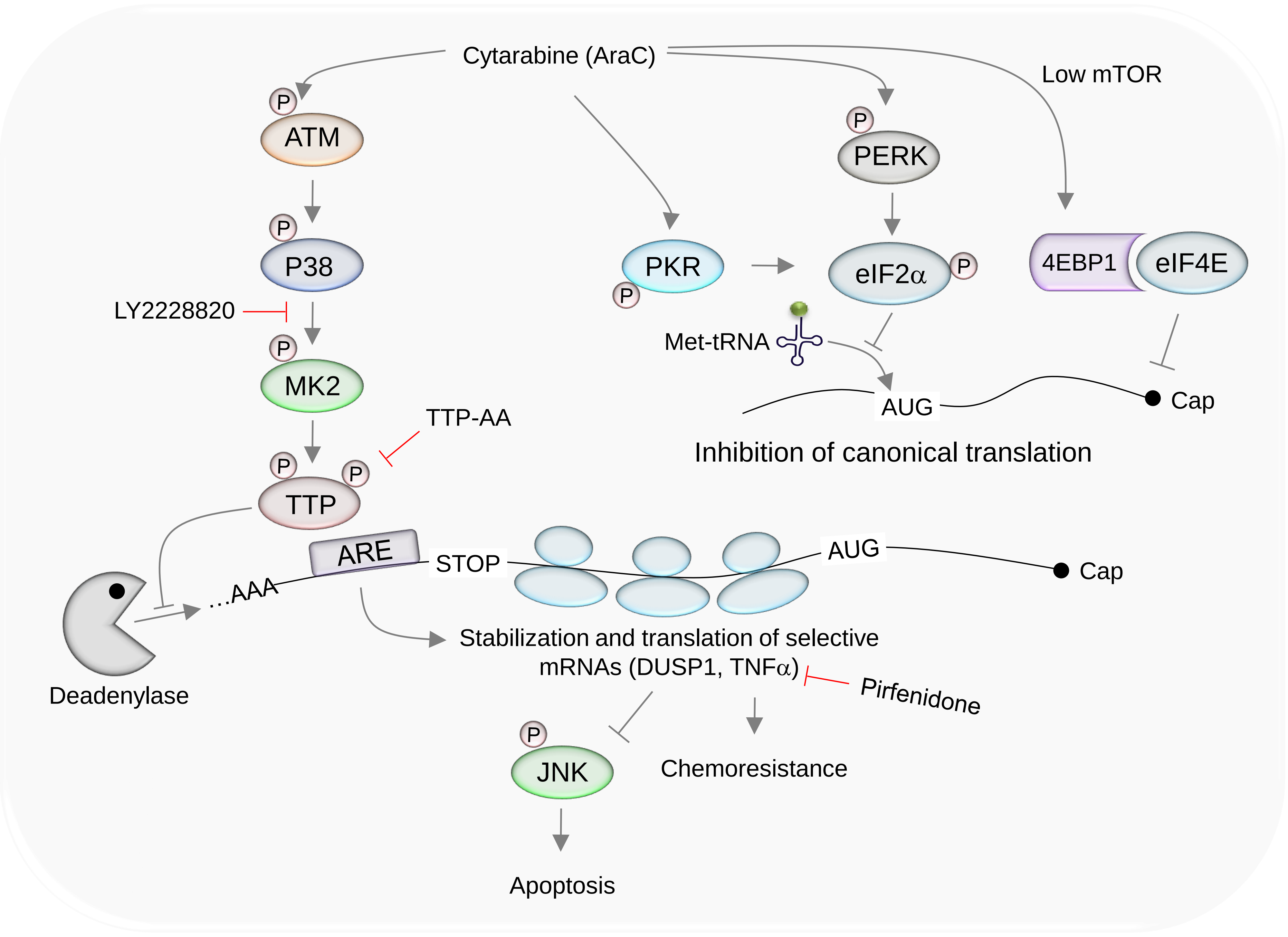
