## Supplemental Information for "A post-transcriptional program of chemoresistance by AU-rich elements and TTP"

**List of Supplemental Information**

Methods (related to main text, main figures 1-8 & supplemental figures S1-S6)

References (related to supplemental information, methods section)

Supplemental Figure legends S1-S6, & Supplemental Table legends, related to main figures 1-8

Supplemental Figures S1-S6 (related to main figures 1-8)

Supplemental Tables S1-2 (related to main figures 1-8)

**Methods**

**Cell Culture**

THP1 cells were cultured in Dulbecco’s modified Eagle medium (RPMI)1460 media supplemented with 10% fetal bovine serum (FBS), 2 mM L-Glutamine, 100 g/mL streptomycin and 100 U/ml penicillin at 37°C in 5% CO_2_. SS THP1 cells were prepared by washing with PBS followed by serum-starvation at a density of 2 × 10^5^ cells/mL and AraCS cells, by treatment with 5 M AraC for 3 days or 9 days. MCF7, HFF, HEPG2 and U2OS cells were cultured in Dulbecco’s modified Eagle medium (DMEM) media with 10% FBS, 2 mM L-Glutamine, 100 g/mL streptomycin and 100 U/ml penicillin, as done previously (1;2). MCF7 cells were serum-starved or treated with 150 uM Doxorubicin. THP1 (TIB-202), MV4:11 (CRL-9591), K562 (CCL243), HFF (SCRC-1041), MCF7 (HTB-22), U2OS (HTB-96) and HEPG2 (HB-8065) were obtained from ATCC. MOLM13 (ACC554), NOMO1 (ACC542) and MONOMAC6 (ACC124) were obtained from DSMZ. Cell lines kindly provided by David Scadden (3) and MOLM13-GFP-Luc by Monica Guzman (4) were tested for Mycoplasma (Promega) and authenticated by the ATCC Cell Authentication Testing Service (3). As previously described (5;6), we used bone marrow derived macrophages (BMDMs) transduced with plasmids coding for doxycycline-inducible GFP–TTP, GFP–TTP-AA or GFP.

**Primary AML patient samples and human monocytes**

All human samples (de-identified) were handled in accordance with IRB protocols to SV (2015P000998/MGH), approved by the Partners Human Research Committee Institutional Review Board /MGH IRB, to DAS, and to TG (DF/HCC 13-583), approved by DF/HCC Office for Human Research Studies. AML samples used in this study were obtained by DAS including: MGH15 - bone marrow 60% blasts, karyotype 46, XX, t(9;11)(p22;q23)[20/20]; MGH22 - peripheral blood, 60% blasts, karyotype 46,XX,t(3;21)(q26;q22),t(9;22)(q34;q11.2)[18]/46,XX[2]; and MGH25 - bone marrow, 90% blasts, karyotype 46,XX[20] and by JL-S and TG including bone marrow samples: EQ1899, CI2095, PO2038, LA2053, NC1866, GO1122, CM2164, MV2192, VD2160, XD2101, VL2317, and OA2500. Bone marrow or peripheral blood mononuclear cells were isolated from de novo AML patients by ficoll density gradient centrifugation and cryopreserved with DMSO in a liquid nitrogen tank. Thawed cells were maintained in RPMI media with 10% FBS for several days before drugs treatment and analyses. Human CD34+ monocytes (2M-101) were obtained from Lonza. Primary cells from MLL-AF9, HoxA9-Meis1 mouse models were provided by DS (3). Mouse primary cells were maintained in RPMI media with 10% FBS, 2 mM L-Glutamine, 100 g/mL streptomycin, 100 U/ml penicillin, 5 ng/ml murine IL-3 and 25 ng/ml murine Stem Cell Factor (SCF).

**In vivo AML mouse models**

AML mouse models have been shown to predict therapy response accurately (7). C57BL/6 and NSG were obtained from MGH Cox-7 Gnotobiotic animal facility of the AAALAC-accredited Center for Comparative Medicine and Services at MGH. C57BL/6 or NSG mice were injected intravenously or subcutaneously with HoxA9-Meis1 or MOLM13 cells expressing luciferase (4;8). IVIS imaging system (Perkin Elmer) were used to confirm engraftment of AML cells. Mice were intraperitoneally injected with 200 µl of luciferase substrate D-Luciferin (15 mg/ml) and anesthetized. Images were taken 5 or 10 minutes after D-Luciferin injection. After confirmation of engraftment by IVIS imaging, mice were randomly assigned to two groups and treated with pirfenidone (100 mg/kg, intraperitoneally), LY2228820 (20 mg/kg, intraperitoneally), AraC (30 mg/kg, intraperitoneally) or saline according at indicated combinations and dosages. Tumor volumes were measured by IVIS imaging at indicated time points.

**Polysome profiling with microarray**

Sucrose was dissolved in lysis buffer containing 100 mM KCl, 5 mM MgCl_2_, 100 g/ml cycloheximide, 2 mM DTT and 10 mM Tris-HCl (pH 7.4). Sucrose gradients from 15% to 50% were prepared in ultracentrifuge tubes (Beckman) as previously described (1;9-11). Cells were treated with 100 g/mL cycloheximide at 37°C for 5 minutes before collecting them. Harvested cell were rinsed with ice-cold PBS having 100 g/mL cycloheximide and then were resuspended in lysis buffer with 1% Triton X-100 and 40 U/mL murine (New England Biolabs) for 20 minutes. After centrifugation of cell lysates at 12,000 x g for 20 minutes, supernatants were loaded onto sucrose gradients followed by ultracentrifugation (Beckman Coulter Optima L90) at 34,000 × rpm at 4 °C for 2 hours in the SW40 rotor. Samples were separated by density gradient fractionation system (Teledyne Isco). RNAs were purified by using TRIzol (Invitrogen) from heavy polysome fractions and whole cell lysates. The synthesized cDNA probes from WT Expression Kit (Ambion) were hybridized to Gene Chip Human Transcriptome Array 2.0 (Affymetrix) and analyzed by the Partners Healthcare Center for Personalized Genetic Medicine Microarray and BUMC facilities. Gene ontology analysis for differentially expressed translatome or proteome was conducted by DAVID 6.7 tools (12) (13). Molecular signatures enriched in AraCS or SS were identified by Gene Set Enrichment Analysis (GSEA) (14).

**Plasmids**

TRIPZ plasmids expressing shRNA against human TNF (V2THS_111606), and miR30a primiR sequences used as control (RHS4750), were obtained from Open Biosystems and MGH cancer center, respectively. Stable cell lines were constructed as described by Open Biosystems. The stable cells expressing shRNA against TNF were induced with 1 μg/mL doxycycline at indicated time points to knockdown TNF. Luciferase reporters to test ARE expression were previously described (9). Cells were treated with 10 ng/ml recombinant TNF (R&D Systems) to activate NFB pathway. Myc-tagged TTP-AA (15;16) was a gift from Nancy Kedersha and Shawn Lyons from Paul Anderson’s lab.

**MTS assay**

MTS assay, a colorimetric quantification of viable cells was conducted as described by the manufacturer, Promega. A volume of 100 l cells was placed in a 96-well plate after drugs treatment. A volume of 20 l MTS reagent (CellTiter 96® Aqueous Non-Radioactive Cell Proliferation Assay) was added to each well followed by incubation at 37°C for 1 hour. Absorbance was measured at 490 nm by using a microplate reader.

**Caspase 3/7 assay**

After drugs treatment, cell death was measured by using caspase-glo® 3/7 assay kit (Promega) according to the protocol provided by the manufacturer. The equal volume of caspase-glo reagent was added to cells, and samples were gently mixed with pipetting. The plates were incubated at room temperature in the dark for 2 hours. The luminescence of each sample was measured in a luminometer (Turner BioSystems).

**Flow cytometry and cell cycle analysis**

Cell proliferation was determined by flow cytometry of cells labeled with propidium iodide and bromodeoxyuridine (BrdU). The cells were incubated with 10 M BrdU for 90 minutes at 37°C in 5% CO_2_ before harvesting. Collected cells were fixed in ice cold 70% ethanol overnight. Cells were washed in PBS and treated with 2 M HCl for 30 min. Cells were incubated for 1 hour with anti-BrdU antibody conjugated to FITC (eBioscience) in the dark, washed and stained with propidium iodide. Samples were filtered through a nylon mesh filter and cell cycle analysis performed on the flow cytometry (17).

**Western blot analysis**

Cells were collected and resuspended in lysis buffer containing 40 mM Tris-HCl (pH 7.4), 6 mM MgCl_2_, 150 mM NaCl, 0.1% NP-40, 1 mM DTT and protease inhibitors (Roche). Samples containing 80 μg of protein were loaded onto 10% or 12% SDS-PAGE (Bio-Rad), transferred to PVDF membranes and processed for immunoblotting. Antibodies against p27 (06-445) and tubulin (05-829) were obtained from Millipore. Antibodies against HES1 (sc-25392), eIF2α (sc-11386), phospho-4EBP1 (sc-1809) and GFP (sc-9996) were from Santacruz. Antibodies against phospho-ATM (ab81292), phospho-PKR (ab32036), DUSP1 (ab138265) and phospho-IRE1 (ab124945) were from Abcam. Antibodies against phospho-PERK (649401) were from Biolegend. Antibodies again TNFα (3707), phospho-p38 MAPK (4511), phospho-MK2 (3007), phospho-eIF2α (9721), TTP (71632), JNK (9252), phospho-JNK (9251) and 4EBP1 (9452) were from Cell Signaling Technology.

**qPCR**

Total RNA was isolated using TRIzol (Invitrogen) according to the manufacturer’s instructions. The cDNA was synthesized from 1 μg of RNA using M-MuLV Reverse Transcriptase (NEB) and random hexamer primer (Promega). qPCRs were run on LightCycler® 480 Instrument II (Roche) using 2 X SYBR green mix (Bio-rad). The primers used in the qPCR were as follows: mouse TNF-α sense 5′-GCCTCTTCTCATTCCTGCTTG-3′, antisense 5′-CTGATGAGAGGGAGGCCATT-3′; mouse Gapdh sense 5’-CATGGCCTTCCGTGTTCCT-3’, antisense 5’-TGATGTCATCATACTTGGCAGGTT-3’; Dusp1 sense 5’-GGCCAGCTGCTGCAGTTTGAG-3’, antisense 5’-AGGTGCCCCGGTCAAGGACA-3’.

**Apoptosis analysis**

Leukemic cells were treated with indicated drug combinations. Annexin V FITC/PI staining was performed with FITC Annexin V Apoptosis Detection Kit I (BD Pharmingen). Flow cytometry analysis and FlowJo software were used to quantitate the percentages of apoptotic cells.

**Colony forming assay**

After treatment with indicated drug combinations, the same number of cells were plated in methylcellulose-based media with human recombinant cytokines (stem cell technology, MethoCult™ H4435). Number of colonies was quantitated in each plate after 10 days.

**Mass Spectrometry**

Multiplex quantitative proteomics analysis was conducted, as previously (18), from S+, SS and AraC treated THP1 leukemic cells.

**Immunoprecipitation**

Expression of GFP-tagged TTP-AA mutant was induced with 1 μg/ml doxycycline prior to 1 μM AraC treatment in TTP-deficient BMDM cells. The cells were cross-linked with UV 254 nm. Cells were lysed in lysis buffer (20 mM Tris, pH 7.5, 150 mM NaCl, 1 mM EDTA, 1 mM EGTA, 1% Triton X-100, 2.5 mM Sodium pyrophosphate, 1 mM β-glycerophosphate, 1 mM Na3VO4, protease inhibitor, RNase inhibitor). Cell lysates were incubated overnight at 4°C with either IgG control or GFP antibody. Protein G agarose (Santacruz) was used to pull down antibody bound RNA-protein complexes.

**Inhibitors**

Pirfenidone (10 to 300 g/ml (19-22)) was obtained from Chemietek. AraC (1 to 10 M (23;24)), LY2228820 (0.03 to 2 M (25-28)), BIRB796 (BIRB, 5µM (29-33)), and JNK-IN-8(34) were from Selleckchem. KU55933 (10 M (35)), BAY 11-7082 (10 M (36)) and D-luciferin were from Cayman Chemical and Doxorubicin (10 to 500 nM (37)) was from Tocris Bioscience.

**Motif, AREs, RNA binding proteins & ribosome occupancy analysis**

The Multiple Em for Motif Elicitation (MEME) software was used to search for cis-elements enriched in 5' UTR of translationally regulated genes (38). Human 5' UTR sequences were retrieved from UCSC table browser (39). In a discriminative mode, 5' UTR sequences of translationally up- or down-regulated genes were used as the primary sequences and 5' UTR sequences of translationally unchanged genes, the control sequences. Motifs were found in the given strand with 6-30 nt motif width. We compared polysome-associated mRNAs with their total RNA levels in serum-starved and AraCS cells to generate the change in ribosome occupancy (RO)(40-42)—which is the ratio of the level of mRNA that is associated with heavy polysomes compared to the total mRNA level of each gene (Fig. 2F, venn diagram, heat map). ARE Score algorithm (43) was used to assess scores of AU-rich elements quantitatively. The list of RNA binding protein genes were obtained from RBPDB database (44).

**Statistical analyses**

All experiments in every figure used at least 3 biological replicates except for microarray, mass spectrometry, and patient sample data. Each experiment was repeated at least 3 times. No statistical method was used to pre-determine sample size. Sample sizes were estimated on the basis of availability and previous experiments (1;2). No samples were excluded from analyses. P values and statistical tests were conducted for each figure. Statistical analyses were conducted using R or Excel. Two-tailed unpaired t-test or Wilcoxon rank sum test was applied to assess statistical significance. SEM (standard error of mean) values are shown as error bars in all figures. Means were used as center values in box plots. P-values less than 0.05 were indicated with an asterisk. E-values were used for the statistical significance in the motif analysis.

**Supplemental Figures (Related to main figures 1-8)**

**Figure S1. Related to main figure 1. A.** IC50 values of standard anti-leukemic chemotherapy, AraC, in AML cell lines (45). THP1 cell line was selected for this study as it shows strong resistance to AraC. **B.** Flow cytometric profiles of S+, SS and AraCS using BrdU and PI staining. **C.** G0 arrest of serum-starved THP1 is assessed by Western analysis of p27 and Hes1 levels. **D.** Polysome profiles of S+, SS (serum starvation for 4 hours, 1 day, 2 days and 4 days) and AraCS (5 M AraC treatment for 3 days and 9 days) THP1 cells. Heavy polysomes (≥3 ribosomes)-associated mRNAs were analyzed by microarray. 'P/M' indicates polysome to monosome ratios. **E.** Gene ontology analysis of differentially expressed genes at the translatome level in response to serum starvation. The statistical significance of enriched gene ontology categories is shown as a heap map. **F.** Scatter plots (i), principal component analysis (PCA) analysis (ii) and unbiased hierarchical clustering (iii) of the translatomes of cells that were serum-starved for indicated times. **G.** Heatmap and boxplot of the expression of LSC gene signature (46) in AraCS and SS cells.

**Figure S2. Related to main figure 2. A.** Western analysis of p27KIP1 (p27) in S+ and SS cells as a marker for G0/G1 arrest, shows that G0 cells are induced by serum-starvation in a number of cancer cell lines. **B.** G0 arrest of serum-starved MCF7 is assessed by Western analysis of p27 and Hes1 levels. **C**. Flow cytometric analysis of S+ and SS cells from MCF7 using BrdU and PI staining. **D.** Polysome profiles of S+ and SS cells from MCF7, U2OS, HepG2 and non-cancerous HFF fibroblasts cell lines. Heavy polysomes (≥ 3 ribosomes) were analyzed by microarray. **E.** GSEA shows gene categories commonly up or downregulated in G0 cells from five different cell lines. Heatmap of normalized enrichment score (NES) is shown. 'N' marks the limited resolution of the proteome in the GSEA. **F.** PCA analysis (i) and unbiased hierarchical clustering (ii) of the translatomes of SS or AraCS cells from five different cell lines are shown. **G**. Heatmap and boxplot of the expression of SASP signature genes in SS and AraCS THP1 cells. **H**. Heatmap of the expression of ER-stress related genes in SS and AraCS cells. **I**. Bar graphs showing the expression of CD47 and HLA-G in S+, SS and AraCS cells. **P* ≤ 0.05 Data are represented as average ± SEM.

**Figure S3. Related to main Figure 3.** **A.** Distinct motifs enriched in 5' UTRs of genes where ribosome occupancy is significantly increased (∆RO↑, top panel) or decreased (∆RO↓, bottom panel) in G0 chemoresistant cells. **B**. Minimum free energy of RNA secondary structure and length of 5'UTRs of genes. **C**. Expression of genes involved in the decay of ARE mRNAs in SS THP1 (top) or SS MCF7 (bottom) compared to S+ cells is shown by GSEA. **D.** Heatmap of the expression of exosome complex genes (3’-5’ exonuclease RNA decay and processing complex) in G0 cancer cells. **E.** Boxplot showing reduced expression of proteasome complex genes in G0 leukemic cells. **F.** Expression of ARE-binding proteins in G0 leukemic cells are shown as a heatmap. These proteins are known to cause ARE mRNA decay or translation repression. Data are represented as average ± SEM.

**Figure S4. Related to main Figures 4-6. A**. Western analysis of indicated proteins in THP1 cells at time points after serum starvation or 5 µM AraC treatment. **B**. Relative ratio of phospho-p38 MAPK to total p38 MAPK, phospho-MK2 to total MK2, or TTP to tubulin (loading control), at indicated time points upon AraC or SS treatment. **C**. Effect of p38 MAPK inhibition on survival of AraC-resistant cells. MOLM13 leukemic cells were pre-treated with various concentrations of BIRB796 (BB), followed by AraC or vehicle treatment. Cell viability and death were assessed by cell counting, MTS and caspase 3/7 assays. **D.** Flow cytometric profiles of MV4:11 or MOLM13 cells treated with AraC or AraC plus LY2228820. Percentages of BrdU-positive cells are shown. **P* ≤ 0.05 Data are represented as average ± SEM.

**Figure S5. Related to main Figures 5-7. A.** TNF-R1 and TNF-R2 expression at the translatome level. **B**. Translatome expression of BCL2A1, BCL3 and BCL6 at indicated time points after serum starvation. **C**. Flow cytometric profiles of THP1 cells treated with vehicle or recombinant TNF. **D**. Effect of PFD on survival of AraC-resistant cells. THP1 cells pre-treated with 300 µg/ml PFD for indicated time followed by 5 µM AraC. Cell viability and death were measured. **E.** MOLM13-GFP-luciferase cells were cultured with or without HS-27 cells and treated with PLA therapy or AraC. Representative microscopic images (top panel) and quantification of luciferase activity (bottom panel) are shown. **F**. TNFα expression at the translatome (right) and RNA levels (left) in SS cells treated with vehicle or PFD. **G**. Kaplan-Meier survival curves of C57BL/6 mice engrafted with primary HOXA9-Meis1 cells and treated with PLA therapy or AraC in figure 8F. **H.** MCF7 cells were pre-treated with 300 g/ml PFD or vehicle 1 day before treatment with serum starvation or 150 nM doxorubicin. Cell viability of PFD-treated compared to vehicle-treated cells is shown. **P* ≤ 0.05 Data are represented as average ± SEM.

**Figure S6. Related to main Figures 1-8.** A model for chemoresistance in AML. Post-transcriptional and translational regulation of gene expression leads to chemoresistance and G0 cell survival, and is regulated by DNA damage and stress signaling, triggered in subpopulations of cancer by chemotherapy and serum-starvation. AraC or SS treatment induce**s** and enriches for quiescent, chemoresistant leukemic cells. Even though canonical translation is inhibited, ARE-bearing mRNAs are increased and highly translated in G0 cells. Mechanistically, the p38 MAPK-MK2 pathway stabilizes ARE mRNAs such as TNF and DUSP1 via phosphorylation of TTP. DUSP1 inhibits JNK-mediated apoptosis and TNF increases anti-apoptotic genes and cell survival, leading to chemoresistance. Inhibition of ARE mRNA expression by p38 MAPK and TNFα inhibitors or TTP-AA mutant sensitizes resistant leukemic cells to AraC treatment.

**Supplemental Table S1.** **Related to main Figures 1-8.** Genes up-regulated at the translatome level in both AraC and SS cells.

**Supplemental Table S2. Related to main Figures 1-8.** Genes bearing AU-rich elements (AREs) and up-regulated at the translatome level.
